# A druggable ATP13A3–antizyme switch controls adaptive polyamine uptake in cancer

**DOI:** 10.64898/2026.07.01.735759

**Authors:** Arthur Schoofs, Jialin Chen, Rania Abou El Asrar, Eline Ryckaert, Lennert Delarbre, Mujahid Azfar, Natalie Gia-Hue Lũ, Arti Sukhai, Marijke De Jaeger, Stephanie Vrijsen, Suravi Chakrabarty, Jolien Bejster, Emily Meeus, Youri Fayt, Anna Vercauteren, Elke Ausloos, Chris Van den Haute, Rik Gijsbers, Patrizia Agostinis, Steven Verhelst, Noriyuki Murai, Jan Eggermont, Sarah van Veen, Peter Vangheluwe

## Abstract

Cellular polyamine depletion is a promising anticancer strategy, but compensatory polyamine uptake limits efficacy when synthesis is blocked by DFMO (difluoromethylornithine), a clinically approved inhibitor of ornithine decarboxylase. The transporter and feedback mechanism driving this adaptive response have remained unclear. Despite their similar biochemical properties, we identify ATP13A3, rather than ATP13A2, as the principal DFMO-responsive polyamine importer, suggesting that these isoforms regulate distinct polyamine fluxes. Mechanistically, the polyamine sensor antizyme not only restrains polyamine biosynthesis but also selectively inhibits ATP13A3-mediated uptake, a brake that is relieved upon DFMO treatment. This regulatory circuit exposes distinct polyamine-acquisition states across cancers, defining synthesis- and/or uptake-biased subtypes that can shift during disease progression. Melanoma metastasis and vemurafenib resistance evolve toward increased ATP13A3-dependent uptake. The polyamine uptake branch controlled by ATP13A3-antizyme regulation can be pharmacologically blocked by AMXT 1501, which directly inhibits ATP13A3. Together, our findings explain DFMO adaptation through ATP13A3–antizyme control and establish ATP13A3 as a targetable node for polyamine depletion strategies in multiple cancers, supporting ongoing clinical evaluation of combined DFMO/AMXT 1501 therapy.

## Introduction

Polyamines such as putrescine, spermidine, and spermine are essential metabolites that among other processes support cell proliferation, translation, and chromatin organization (*1*). However, their intracellular levels must be tightly constrained: polyamine depletion limits growth and survival, whereas excessive accumulation becomes toxic (*2*). Consequently, polyamine homeostasis is controlled by sophisticated regulatory feedback mechanisms, and its disruption contributes to diverse pathologies ranging from cancer to neurodegeneration (*3*).

Cells acquire polyamines through two major inputs, *i.e. de novo* biosynthesis and uptake from the extracellular environment (**Fig. 1A**) (*4*). Biosynthesis is initiated by ornithine decarboxylase 1 (ODC1), the rate-limiting enzyme that converts ornithine into putrescine, whereas uptake occurs through the polyamine transport system (*3*). These two arms of polyamine homeostasis are coordinated by antizyme, a polyamine-induced negative regulator that is expressed when intracellular polyamine levels rise (*5, 6*). Members of the antizyme family (AZ1–3; *OAZ1–3*) bind ODC1 and promote its proteasomal degradation, thereby suppressing polyamine biosynthesis in a negative feedback loop (*7*). Remarkably, antizyme has also been known for decades to inhibit cellular polyamine uptake (*8*), but the transporter(s) through which antizyme regulates polyamine uptake has (have) remained unknown. This missing link has prevented a mechanistic understanding of how cells coordinate polyamine synthesis and import to maintain homeostasis.

**Figure 1:**
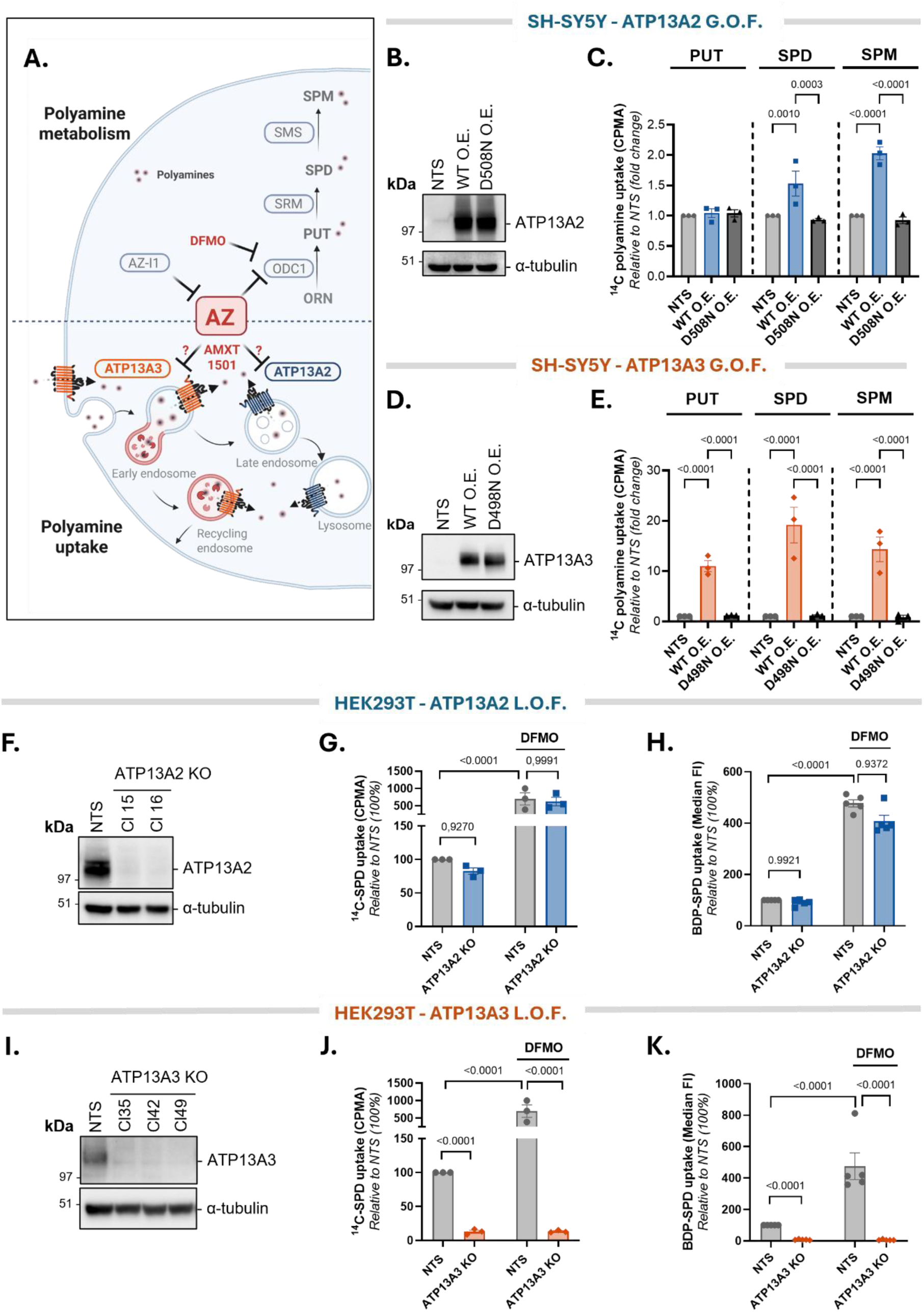
ATP13A3 is the dominant transporter in cellular polyamine uptake that responds to DFMO. Comparative analysis of ATP13A2- and ATP13A3-mediated polyamine uptake in SH-SY5Y gain-of-function (G.O.F.) overexpression models and HEK293T loss-of-function (L.O.F.) knockout (KO) models. **(A)** Schematic overview of intracellular polyamine metabolism and uptake, including inhibition of ornithine decarboxylase 1 (ODC) by α-difluoromethylornithine (DFMO) and the proposed subcellular localization of ATP13A2 (late endo/lysosomes) and ATP13A3 (early/recycling endosomes, plasma membrane). Question marks indicate the unknown target of antizyme (AZ) and the polyamine transport inhibitor AMXT 1501. Abbreviations: AcSPD, N¹-acetylspermidine; AcSPM, N¹-acetylspermine; AZIN, antizyme inhibitor; dcSAM, decarboxylated S-adenosylmethionine; ER, endoplasmic reticulum; ORN, ornithine; PAOX, polyamine oxidase; PUT, putrescine; SAM, S-adenosylmethionine; SAT1, spermidine/spermine N¹-acetyltransferase 1; SMOX, spermine oxidase; SMS, spermine synthase; SPD, spermidine; SPM, spermine; SRM, spermidine synthase. **(B**, **D)** Representative immunoblots showing ATP13A2 (**B**) or ATP13A3 (**D**) protein levels in SH-SY5Y cells stably overexpressing wild-type (WT) ATP13A2 or ATP13A3 or the corresponding catalytically inactive D508N or D498N mutant, respectively. Non-transduced (NTS) cells served as controls, and α-tubulin was used as a loading control. Blots are representative of three independent biological replicates. **(C, E)** Uptake of ¹⁴C-radiolabeled PUT, SPD, and SPM (5 µM, 30 min) in the corresponding ATP13A2 **(C)** or ATP13A3 **(E)** overexpression models. Radiolabelled uptake was quantified by counts per minute (CPMA). Data are presented as uptake normalised to the NTS control for each polyamine, with individual biological replicate values shown as dots and bars representing the mean ± SEM of *n* = 3. Two-way ANOVA followed by Tukey post hoc multiple comparison test, was performed on these data. **(F**, **I)** Immunoblot confirmation of ATP13A2 (**F**) and ATP13A3 (**I**) KO clones (Cl) in HEK293T cells; α-tubulin served as a loading control. Blots are representative of three independent biological replicates. **(G, H, J, K)** Uptake of 14C-labelled SPD (1 µM, 30 min) and BODIPY-labelled SPD (BDP-SPD; 0.1 µM, 30 min) in HEK293T NTS cells, ATP13A2 **(G, H)** and ATP13A3 **(J, K)** KO cells in the presence or absence of pre-treatment with 1.5 mM DFMO for 24h. Radiolabelled uptake was quantified by counts per minute (CPMA) and BDP-SPD uptake was quantified as median fluorescence intensity. Data are presented as uptake normalised to untreated NTS (set to 100%), with individual biological replicate values shown as dots and bars representing the mean ± SEM of *n* = 3 or 5. Statistical analyses were performed on log₁₀-transformed raw uptake values using two-way ANOVA followed by Šidák’s post hoc multiple comparisons test. Normality was assessed using a QQ plot.

Resolving this question is particularly important in cancer. Elevated polyamine pools support the rapid growth and proliferation of cancer cells (*4*). This dependency has motivated therapeutic strategies aimed at depleting intracellular polyamines, most notably through inhibition of ODC1 using difluoromethylornithine (DFMO, eflornithine) (*9, 10*). DFMO has reached clinical use for maintenance therapy in high-risk neuroblastoma (*11*) and is being explored more broadly to achieve polyamine-depletion strategies in cancer (*12–15*). However, a major limitation of DFMO-based therapy is cellular adaptation. When polyamine synthesis is inhibited, cells can compensate by increasing polyamine uptake from their environment (*16, 17*). This adaptive response undermines therapeutic efficacy, which has stimulated the development of polyamine transport inhibitors (*10, 18, 19*), which have shown promise in combination with DFMO in preclinical models of neuroblastoma (*12*), glioblastoma (*13, 14, 18*) and pancreatic cancer (*13*). Among these, the polyamine analog AMXT 1501 is currently being evaluated in combination with DFMO in clinical trials (NCT07287917, NCT06465199, NCT05717153), which in Phase I appeared safe and tolerated with evidence of preliminary clinical activity (NCT03536728) (*15*). Yet the molecular mechanism underlying DFMO-induced uptake and the transporters targeted by AMXT 1501 and its mode of action remain poorly understood.

Our recent work has established the P5B-type transport ATPases as central components of the mammalian polyamine transport system (*3*). ATP13A2 (*20, 21*), ATP13A3 (*21–24*), and ATP13A4 (*25*) have all been implicated in cellular polyamine uptake, whereas ATP13A5 remains unexplored. Intriguingly, these transporters are associated with strikingly different human diseases. ATP13A2 dysfunction causes neurodegenerative disorders including Kufor-Rakeb syndrome and Parkinsonism (*26, 27*), whereas ATP13A3 loss-of-function is linked to pulmonary arterial hypertension (*28, 29*), and ATP13A4 has been associated with neurodevelopmental disorders (*25*). The existence of multiple disease-associated polyamine transporters therefore presents a fundamental paradox. If these proteins all mediate polyamine uptake, why are their physiological and pathological consequences so distinct? Whether there is transporter hierarchy in the regulation of polyamine uptake and/or whether these transporters represent differentially regulated components of a common uptake system remains unclear. ATP13A2 and ATP13A3 represent the two ubiquitous isoforms, whereas ATP13A4 and ATP13A5 exhibit restricted tissue distribution (*3*). Recent work provided important clues that ATP13A3 may play a central role: ATP13A3 is implicated in the compensatory polyamine uptake response induced by DFMO in neuroblastoma cells (*16*), whereas ATP13A3 expression has been independently associated with worse prognosis in neuroblastoma (*16*), pancreatic (*23*), head and neck (*30*), and breast cancer (*31*). These observations raise the broader questions whether ATP13A3 represents the main polyamine uptake route in mammalian cells under basal and/or DFMO conditions, and whether it is mechanistically coupled to polyamine synthesis via antizyme control.

Here, we firmly establish ATP13A3 as the key cellular polyamine importer that falls under control of antizyme and DFMO, demonstrating that ATP13A3 fulfills a central role in polyamine homeostasis that complements polyamine synthesis. This mechanism explains DFMO-induced compensatory uptake in cancer, which can be prevented by AMXT 1501 that directly inhibits ATP13A3. Finally, dysregulation of the ATP13A3-antizyme pathway emerges across multiple cancer types and is reinforced during melanoma metastasis and vemurafenib resistance, identifying ATP13A3 as a druggable target in cancers that escape polyamine homeostatic control.

## Results

### ATP13A3 represents the dominant DFMO-responsive polyamine uptake route

The two ubiquitous polyamine transporters ATP13A2 and ATP13A3 have previously been implicated in cellular polyamine uptake (*16, 20–24*), but their relative impact has not yet been accurately assessed. We therefore directly compared ATP13A2 and ATP13A3-mediated polyamine uptake in matched gain- and loss-of-function cell systems.

First, we generated SH-SY5Y human neuroblastoma cells stably expressing wild-type (WT) ATP13A2, catalytically inactive ATP13A2-D508N, WT ATP13A3, or catalytically inactive ATP13A3-D498N (**Fig. 1B,D)**. Protein quantification against protein standards with purified ATP13A2 or ATP13A3 (**Fig. S1; Fig. S2A-B**) illustrated that ATP13A2 was expressed at more than 2-fold higher levels than ATP13A3 (**Fig. S2C-D**). However, ATP13A2 caused only a modest increase in radiolabeled polyamine uptake, enhancing 14C-spermidine and 14C-spermine uptake by approximately 1.5- and 2-fold, respectively, while leaving 14C-putrescine uptake unchanged. This activity was absent for ATP13A2-D508N, confirming dependence on the transport cycle (**Fig. 1C**). Under the same conditions, ATP13A3 had a much stronger effect increasing uptake of all three major polyamines, with approximately 10-fold higher 14C-putrescine, 20-fold higher 14C-spermidine, and 15-fold higher 14C-spermine uptake. The catalytically inactive ATP13A3-D498N mutant failed to enhance uptake (**Fig. 1E**). Thus, although ATP13A2 can contribute to uptake of selected polyamines, ATP13A3 is markedly more effective at driving cellular polyamine import and displays a broader polyamine uptake profile. We rule out that polyamine degradation in serum plays a role, since experiments were performed with heat-inactivated fetal calf serum **(Fig. 1C, E)** or supplementation of aminoguanidine (**Fig. S3A,B**) to kill serum polyamine oxidase activity.

We next complemented our analysis with HEK293T knockout (KO) cells of ATP13A2 or ATP13A3 to assess their relative endogenous contribution on cellular polyamine uptake. After validating the isoform-specific KO by immunoblotting (**Fig. 1F, I**), we measured polyamine uptake side by side with radiolabeled polyamines (**Fig. 1G,J; Fig. S3C-F**) as well as with BODIPY-conjugated fluorescent polyamine analogs (**Fig. 1H,K; Fig. S3G-J**), which have been validated as transported substrates of the P5B-type polyamine transporters ATP13A2 and ATP13A3 (*21, 32*). Surprisingly, loss of ATP13A2 caused only a minor, non-significant 5-20% decrease in basal cellular polyamine uptake (**Fig. 1G,H; Fig. S3C,D,G,H**). However, the impact of ATP13A3 KO was much more prominent with a robust reduction in uptake of 14C-putrescine, 14C-spermidine and 14C-spermine uptake of around 90% (**Fig. 1J; Fig. S3E,F**). We also observed a 90% and 60% significant decrease of BODIPY-spermidine and BODIPY-spermine uptake, albeit a non-significant decrease in BODIPY-putrescine (**Fig. 1K; Fig. S3I,J**). This further illustrates that ATP13A3 is a major contributor to polyamine uptake in cells (*16, 22, 23*).

We then asked which transporter mediates the adaptive uptake response to polyamine depletion following DFMO treatment. Consistent with compensatory activation of the polyamine transport system after ODC1 inhibition (*16, 17*), DFMO significantly increased cellular uptake of all radiolabeled polyamine species with a factor between 4- and 7-fold (**Fig. 1G,J; Fig. S3C-F**), and a 4-fold increase in BODIPY-spermidine (**Fig. 1H,K**) (which was less pronounced with BODIPY-putrescine and BODIPY-spermine **(Fig. S3G-J**). Strikingly, the DFMO-induced increase in polyamine uptake was completely abolished by ATP13A3 KO (**Fig. 1J,K; Fig. S3E,F,I,J**), whereas ATP13A2 loss had no significant impact (**Fig. 1G-H; Fig. S3C,D,G,H**). Because cellular uptake of radiolabeled polyamines and BODIPY-spermidine are similar under basal and DFMO conditions, we decided to monitor BODIPY-spermidine uptake to further investigate the DFMO-mechanism. Thus, ATP13A3 is the main contributor to basal cellular polyamine uptake and mediates the increased polyamine import induced by DFMO, establishing a functional hierarchy among P5B-type polyamine transporters in the regulation of cellular polyamine uptake.

### ATP13A3 is a bona fide polyamine-transporting P5B-type ATPase

Our work positions ATP13A3 as a central mediator of adaptive polyamine uptake in response to polyamine depletion. Despite its major role in polyamine uptake, ATP13A3 has not been biochemically established as a polyamine transporter. We therefore determined the biochemical properties and polyamine selectivity of purified human ATP13A3 protein and compared it with ATP13A2, which we have previously characterized (*20*). Human ATP13A3 shares 41.1% sequence identity and 58.7% sequence similarity with ATP13A2 (**Fig. S4A**) and is predicted to adopt a similar P5B-type ATPase architecture as ATP13A2 (*33–35*), comprising an N-terminal domain (NTD), actuator domain (A), nucleotide-binding domain (N), phosphorylation domain (P), and membrane domain (M) (AlphaFold3 models in **Fig. 2A**; **Fig. S4B**) (*36*). Within the M domain, ATP13A3 contains all residues to shape an extracytosolic-facing, polyamine-binding pocket (**Fig. 2B; Fig. S4C**) that has been described for ATP13A2 in cryo-EM structures (*33–35*).

**Figure 2:**
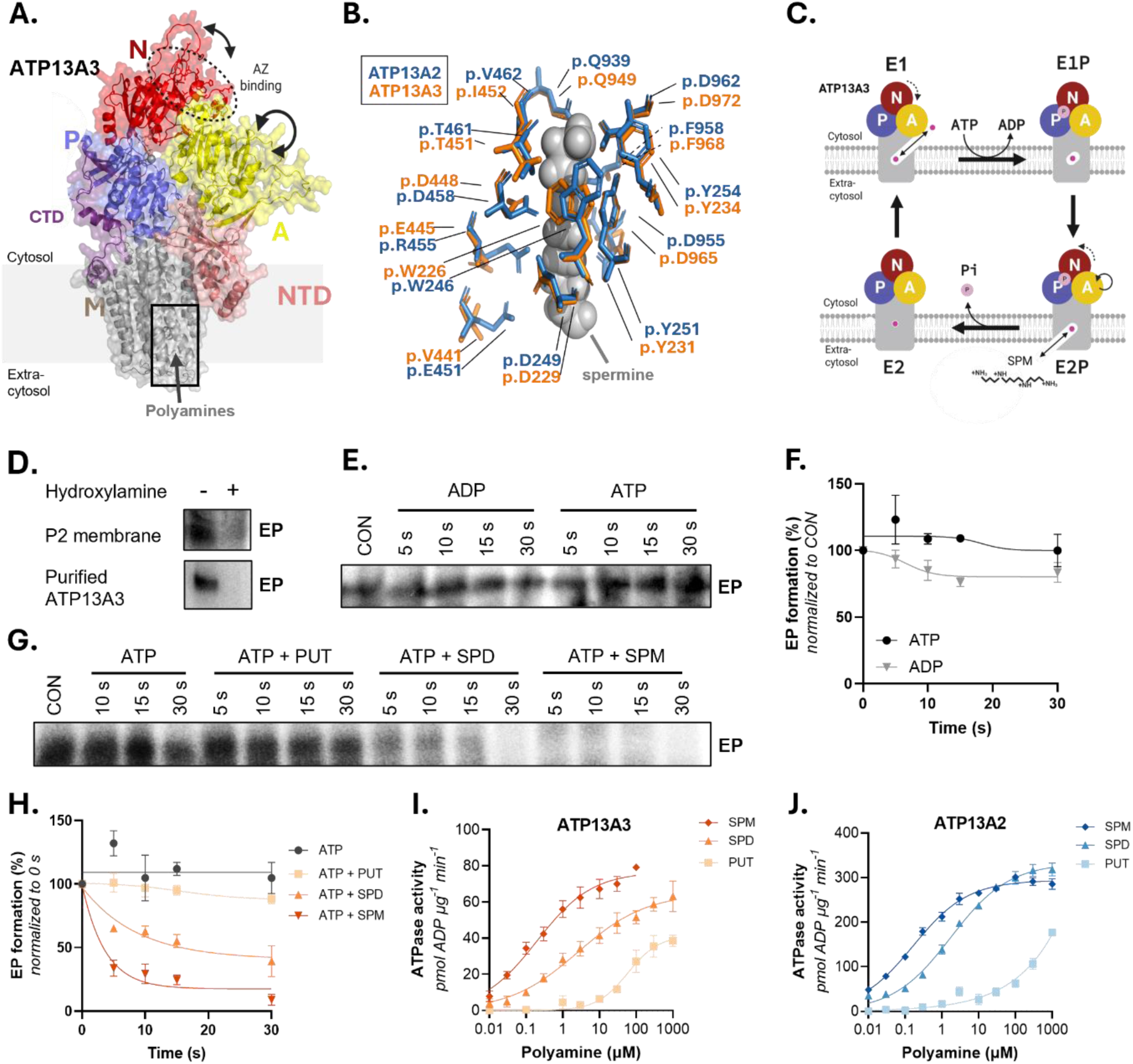
ATP13A3 is a P5B-type polyamine transporter with ATP13A2-like properties. Biochemical characterization of purified human ATP13A3 by autophosphorylation and ATPase assays. **(A)** AlphaFold structural model of ATP13A3 displaying the actuator (A), phosphorylation (P), nucleotide-binding (N), membrane (M) and N-terminal domains (NTD). The proposed antizyme (AZ)-binding region identified in this study is indicated by a dashed oval. The conserved polyamine binding site is indicated by a box that is shown in panel B. The arrow indicates the polyamine transport direction from the extracytosolic (*i.e.* extracellular and/or endosomal lumen) to the cytosolic side. **(B)** Structural overlay of the ATP13A3 (AlphaFold 3prediction; orange) and ATP13A2 (cryo-EM structure of ATP13A2 in the spermine bound E2-Pi state. PDB: 7n78 (72); blue) polyamine-binding pockets, highlighting the conserved residues shaping the substrate pocket. Spermine is shown as grey spheres. **(C)** Schematic representation of the Post-Albers catalytic cycle of ATP13A3, based on the established transport mechanism of the closely related P5B ATPase ATP13A2, illustrating the E1, E1P, E2P, and E2 conformational states. ATP binding and phosphorylation generate the E1P intermediate, whereas polyamine substrate binding promotes dephosphorylation of the E2P intermediate. **(D)** Detection of the ATP13A3 phosphoenzyme (EP) intermediate in human ATP13A3-expressing P2 membrane fractions isolated from *Saccharomyces cerevisiae* and in human ATP13A3 purified from HEK293T cells. Hydroxylamine (0.3 M) sensitivity confirms the formation of an aspartyl-phosphate intermediate characteristic of P-type ATPases. Representative radiogram of *n* = 3 independent biological replicates. **(E-F)** Representative autoradiogram **(E)** and quantification **(F)** of pulse ([γ-³²P]-ATP) chase (cold ATP or ADP) experiments to determine the ATP and ADP sensitivity of the ATP13A3 EP intermediate. Data are presented as mean ± SEM of *n* = 4 independent biological replicates. **(G-H)** Representative autoradiogram **(G)** and quantification **(H)** of pulse ([γ-³²P]-ATP) chase (cold ATP) experiments in the presence of 1 mM putrescine (PUT), spermidine (SPD), or spermine (SPM) to assess substrate-induced dephosphorylation of ATP13A3. Data are presented as mean ± SEM of *n* = 3 independent biological replicates. **(I-J)** Dose-response curves showing the effect of PUT, SPD, and SPM on the ATPase activity of human ATP13A3 (**I**) and ATP13A2 (**J**) purified from HEK293T cells. Data are presented as mean ± SEM of *n* = 4 independent biological replicates.

Next, we examined the catalytic hallmarks of the purified human ATP13A3 transporter from two independent expression systems, *i.e. Saccharomyces cerevisiae* and HEK293T cells (**Fig. S1**). In line with the P-type ATPase catalytical cycle (**Fig. 2C**), ATP13A3 formed a phosphoenzyme intermediate (EP) upon incubation with radiolabeled ATP, both in membrane fractions as in purified detergent-solubilized preparations (**Fig. 2D**), just as ATP13A2 (*20*). This intermediate was hydroxylamine-sensitive, consistent with an aspartyl autophosphorylation on the catalytic aspartate in the conserved 498DKTG motif of the P-domain (*20*). ATP13A3 phosphoenzyme levels were only weakly sensitive to ADP and did not respond to ATP chase, indicating that ATP13A3 does not efficiently complete its catalytic cycle and is accumulating in the E2P state (**Fig. 2E-F**). Addition of the transported substrate triggered phospho-enzyme turnover (**Fig. 2G-H, Fig. S4D-I**). Spermidine and spermine accelerated phosphoenzyme dephosphorylation in a concentration-(**Fig. S4D-E**) and time-dependent (**Fig. 2G-H**) manner, whereas putrescine had little effect under these low-temperature phosphoenzyme assay conditions (**Fig. 2G-H, S4D-I**). The same substrate-dependent dephosphorylation profile was observed with ATP13A3 purified from yeast or HEK293T cells, excluding expression-system effects (**Fig. S4F-I**).

We directly compared the ATP13A3 and ATP13A2 substrate preference in an independent ATPase assay at 37°C. Spermine, spermidine and putrescine stimulated ATP13A3 ATP hydrolysis in a concentration-dependent manner with markedly different apparent affinities. Spermine activated ATP13A3 most potently, followed by spermidine (**Fig. 2I**), similar to ATP13A2 (**Fig. 2J**) (**Table 1**). ATP13A2 and ATP13A3 only respond to high concentrations of putrescine with ATP13A3 presenting a higher apparent affinity to putrescine than ATP13A2 (**Fig. 2I-J**) (**Table 1**), which is in line with the polyamine selectivity in cellular uptake (**Fig. 1C,E; Fig. S3C-F**). Under the same experimental conditions, ATP13A2 presents a higher maximal turnover as compared to ATP13A3 (**Fig. 2I-J**)(**Table 1**) although the maximal turnover is dependent on the detergent/lipid composition. ATP13A2 turnover is regulated by phospholipids such as phosphatidic acid (PA) and phosphatidylinositol(3,5)bisphosphate (PI(3,5)P₂) (*20, 37*), and we examined lipid regulation of ATP13A3 activity. First, we performed protein-lipid overlays using a purified recombinant N-terminal protein fragment of ATP13A3 to select candidate regulatory lipids (**Fig. S5A-D**), as previously done for ATP13A2 (*37*). The functional effect of identified candidate lipids was then assessed on the ATP13A3 ATPase activity in the reference background of phosphatidylcholine (PC) re-lipidation, which stabilized ATP13A3 activity (**Fig. S5E**). Compared to PC, the candidate regulatory lipids, phosphatidylinositol(3)phosphate (PI(3)P), PI(3,5)P2 and sulfatide further stimulated ATP13A3 activity, whereas phosphatidic acid (PA) had opposite effects (**Fig. S5F**), demonstrating that the maximal turnover of ATP13A3 is dynamically regulated by the lipid composition.

**Table 1.** Apparent kinetic parameters of ATP13A3 and ATP13A2.

|  | ATP13A3 |  |  | ATP13A2 |  |  |
| --- | --- | --- | --- | --- | --- | --- |
|  | V<sub>max</sub><br>(pmol ADP μg<sup>-1</sup> min<sup>-1</sup>) | n<br>(Hill coefficient) | K<sub>m</sub><br>(μM) | V<sub>max</sub><br>(pmol ADP μg<sup>-1</sup> min<sup>-1</sup>) | n<br>(Hill coefficient) | K<sub>m</sub><br>(μM) |
| **SPM** | 76.7 ± 3.8 | 0.58 ± 0.09 | 0.20 ± 0.06 | 294.1 ± 4.7 | 0.58 ± 0.04 | 0.17 ± 0.02 |
| **SPD** | 64.8 ± 5.2 | 0.46 ± 0.08 | 2.61 ± 1.39 | 332.6 ± 8.6 | 0.55 ± 0.04 | 1.76 ± 0.29 |
| **PUT** | 41.5 ± 4.4 | 1.02 ± 0.26 | 58.92 ± 20.62 | N.D. (>200) | 0.38 ± 0.10 | N.D. (>200) |

Apparent kinetic parameters derived from the ATPase activity measurements of purified human ATP13A3 and ATP13A2 isolated from HEK293T cells. ATPase activities were fitted using Hill function. Values are reported as mean ± SEM.

Together, our biochemical data establish ATP13A3 as a bona fide P5B-type polyamine transporter with remarkably similar substrate preferences, biochemical properties and overlapping lipid regulation as compared to ATP13A2.

### Antizyme selectively binds and inhibits ATP13A3

The dominant role of ATP13A3 in cellular and DFMO-induced polyamine uptake makes ATP13A3 an exquisite candidate for antizyme regulation. This is supported by AlphaFold3 predictions that indicate preferential interaction of antizyme isoforms with ATP13A3 rather than ATP13A2 (*36*). The predicted ATP13A3 complexes with AZ1, AZ2, and AZ3 showed interface confidence scores (iPTM) in the 0.6–0.8 range (medium to high confidence), whereas predicted ATP13A2-antizyme complexes scored below the iPTM threshold of 0.6 (low confidence) (**Fig. 3A-B, Fig. S6A-B**). This distinction is further supported by the predicted aligned error matrices (PAE) (**Fig. S6C-H**) for ATP13A2 and ATP13A3 complexes with AZ1-3. AlphaFold3 models predicted all antizyme isoforms to bind on the same position on the cytosolic headpiece of ATP13A3 (**Fig. 3A-B, Fig. S6B, Fig. S7A**), despite considerable sequence variations among antizyme isoforms (36% to 56% sequence identity **Fig. S6I**). The ATP13A3 and AZ2 interaction interfaces display similar interaction profiles (**Fig. S7B,D**), whereas the ATP13A3 residues, predicted to interact with AZ2, are not conserved in ATP13A2 (**Fig. S7C**), suggesting that differences in residue composition may account for preferential binding of AZ2 to ATP13A3.

**Figure 3:**
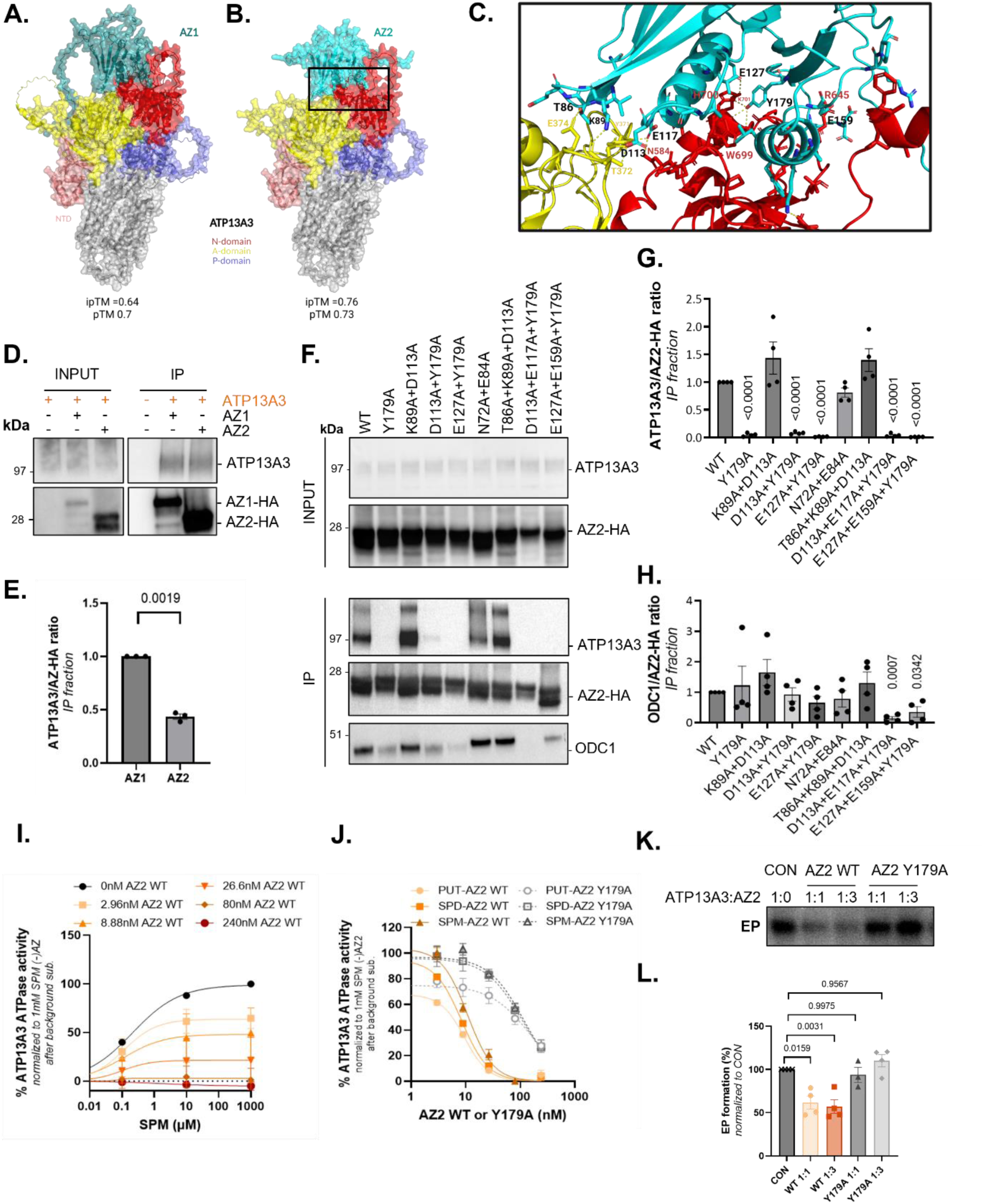
Antizyme 2 selectively binds ATP13A3 and inhibits its catalytic activity. **(A-B)** AlphaFold heterodimer models of ATP13A3 in complex with antizyme 1 (AZ1) **(A)** or antizyme 2 (AZ2) **(B)**. ATP13A3 domains are coloured as follows: N-domain (red), P-domain (purple), A-domain (yellow), transmembrane domain (grey), and N-terminal domain (NTD, pink). AZ1 and AZ2 are shown in cyan. Residues at the predicted ATP13A3-antizyme interaction interface are shown as sticks. Model confidence scores (ipTM and pTM) are indicated. **(C)** Close-up view of the predicted ATP13A3-AZ2 interaction interface highlighting residues selected for mutagenesis. **(D-E)** Co-immunoprecipitation of ATP13A3 with AZ1-HA or AZ2-HA in HEK293T cells. **(D)** Representative immunoblots of input and immunoprecipitated (IP) fractions. **(E)** Quantification of ATP13A3:AZ-HA interaction normalized to AZ1 binding. Data are presented as mean ± SEM of *n* = 3 independent biological replicates, with individual data points shown. Statistical analysis was performed using a two-tailed one-sample t-test. **(F-H)** Co-immunoprecipitation analysis of ATP13A3 and endogenous ODC1 with AZ2 WT and AZ2 interface mutants. **(F)** Representative immunoblots of input and IP fractions. **(G-H)** Quantification of **(G)** ATP13A3:AZ2-HA and **(H)** ODC1:AZ2-HA interactions normalized to AZ2 WT binding. Data are presented as mean ± SEM of *n* = 4 independent biological replicates, with individual data points shown. Two-tailed one-sample t-test. **(I)** Spermine (SPM)-stimulated ATPase activity of purified ATP13A3 in the presence of increasing concentrations of AZ2 WT. ATPase activity was normalized to the activity measured at 1 mM SPM in the absence of AZ2. Data are presented as mean ± SEM of *n* = 4 independent biological replicates. **(J)** Putrescine (PUT), spermidine (SPD), and spermine (SPM)-stimulated ATPase activity of purified human ATP13A3 measured in the presence of increasing concentrations purified human AZ2 WT or AZ2 179A. ATPase activity was normalized to the activity measured in the presence of 1 mM SPM in the absence of AZ2. Data are presented as mean ± SEM of n = 3 independent biological replicates. **(K-L)** Effect of AZ2 on ATP13A3 phosphoenzyme (EP) formation. **(K)** Representative autoradiogram of pulse ([γ-³²P]-ATP) experiments to assess ATP13A3 EP formation (quantified in **L**) in the absence or presence of purified AZ2 WT or the ATP13A3-binding-deficient mutant AZ2 Y179A at ATP13A3:AZ2 molar ratios of 1:1 and 1:3. The radiogram shown here is a cropped image derived from **Fig. S8G**. Data are presented as mean ± SEM of *n* = 4 independent biological replicates, with individual data points shown. Two-tailed one sample t-test.

To experimentally test the predicted ATP13A3-specific interaction, we performed co-immunoprecipitation assays in HEK293T cells with HA-tagged AZ1 and AZ2, the two ubiquitously expressed antizyme isoforms (AZ3 is testis-restricted (*5*)). To eliminate the polyamine-dependent feedback on antizyme expression, AZ1 and AZ2 were expressed from constructs lacking the ribosomal frameshift sequence required for translational regulation. Antizyme co-immunoprecipitations robustly pulled down ATP13A3 (**Fig. 3D**), whereas ATP13A2 showed no detectable interaction under the same conditions (**Fig. S8A**). Quantification indicated stronger ATP13A3 recovery with AZ1 than with AZ2, but both isoforms reproducibly bound ATP13A3 (**Fig. 3E**). Thus, among the two ubiquitously expressed P5B-type polyamine transporters AZ1 and AZ2 selectively interacted with ATP13A3. We next asked whether this interaction directly inhibits ATP13A3 activity. In biochemical assays with purified human ATP13A3 (**Fig. S1C-D**), ATP13A2 (**Fig. S1E-F**) and recombinant AZ2 protein (**Fig. S8B-C**), AZ2 completely blocked polyamine-dependent ATPase activity of ATP13A3 at nanomolar concentrations of AZ2 (IC50 range 9.6-10.8 nM) (**Fig. 3I-J**), but AZ2 did not significantly inhibit ATP13A2 (**Fig. S8F**). AZ2 had a similar impact on putrescine, spermidine and spermine-dependent ATPase activity (**Fig. 3J**). Phosphoenzyme assays further demonstrated that AZ2 reduced the autophosphorylation (**Fig. 3K-L**), but not the substrate-dependent dephosphorylation (**Fig. S8G-H**), which fits well with the predicted interaction site involving the nucleotide binding (N-) domain (**Fig. 3B-C**) that undergoes considerable conformational changes to control autophosphorylation (**Fig. 2C**).

Together, our data conclusively demonstrate a specific functional interaction between ATP13A3 and antizyme with nanomolar affinity.

We further investigated the predicted interface in the AlphaFold3 ATP13A3-AZ2 structural model and selected conserved residues among AZ1 and AZ2 that may contribute to the ATP13A3 interaction (**Fig. S7D-E, H,I**). Via co-immunoprecipitation experiments with several AZ2 variants, we found that AZ2 mutations targeting the ATP13A3 N-domain interface strongly reduced the interaction, whereas mutations in the predicted A-domain interface or outside the interface had little effect (**Fig. 3F-G**). A single AZ2 mutation, Y179A, was sufficient to largely abolish ATP13A3 binding (**Fig. 3F-G**), identifying this residue as a hotspot for the ATP13A3-AZ2 interaction. Importantly, the same residue was required for inhibition, since the inhibitory effect of purified AZ2-Y179A (**Fig. S8D-E**) on ATP13A3 ATPase activity (**Fig. 3J**) and phospho-enzyme formation (**Fig. 3K-L**) was impaired rendering a 10-fold lower IC50 than WT AZ2 (IC50 range 107.2-147.0 nM). Strikingly, the experimentally confirmed binding site on AZ2 for ATP13A3 (**Fig. S7D-F**) also overlaps with the position that the complementary interface on AZ1 uses for ODC1 interaction (**Fig. S7G**) (*7*). Indeed, co-immunoprecipitation of AZ2 not only pulled down ATP13A3, but also ODC1 (**Fig. 3F**). The ODC1 and ATP13A3 interactions were both disturbed by mutations in the same antizyme region, although the single Y179A mutation was insufficient to disturb the ODC1 interaction (**Fig. 3F, H**). This illustrates that ATP13A3 and ODC1 occupy an overlapping interaction site on antizyme, which also corresponds to the established binding site of antizyme inhibitor 1, AZ-I1 (*AZIN1*) the negative regulator of antizyme (**Fig. S7G**) (*7*).

Together, our findings identify ATP13A3 as a direct target for negative control by antizyme. Antizyme engages with ATP13A3 at nanomolar affinity overlapping with the canonical ODC1-binding interface, thereby suppressing polyamine ATPase activity.

### DFMO activates polyamine uptake by releasing ATP13A3 from antizyme control

Having identified ATP13A3 as a direct biochemical target of antizyme, we next asked whether this represents the missing molecular explanation for how antizyme suppresses cellular polyamine uptake under basal and/or DFMO-induced conditions (**Fig. 1J-K**).

We first tested whether antizyme controls ATP13A3-dependent uptake in cells by expressing AZ1 or AZ2 in parental HEK293T cells and in ATP13A3 KO cells (**Fig. 4A**), and measured BODIPY-spermidine uptake under basal conditions (**Fig. 4B**) and after DFMO treatment (**Fig. 4C**). In basal conditions, AZ1 or AZ2 expression lowered polyamine uptake only in the control cells, but not in ATP13A3 KO cells (**Fig. 4B**), confirming the specificity of antizyme towards ATP13A3 in cells. DFMO increased BODIPY-spermidine uptake, consistent with activation of compensatory polyamine import (**Fig. 1J-K**). Expression of either AZ1 or AZ2 blunted this uptake response, whereas AZ1 and AZ2 had no effect in ATP13A3 KO cells after DFMO (**Fig. 4C**). ATP13A3 protein expression was not significantly altered between the conditions, suggesting that it is the activity of ATP13A3 that is regulated by antizyme (**Fig. S9A**). Parental HEK293T cells with ATP13A3 overexpression showed an increased BODIPY-SPD uptake, which was countered by AZ1 or AZ2 expression (**Fig. S9B**). Together, the results established that the antizyme-sensitive component of cellular uptake under basal and DFMO conditions requires ATP13A3.

**Figure 4:**
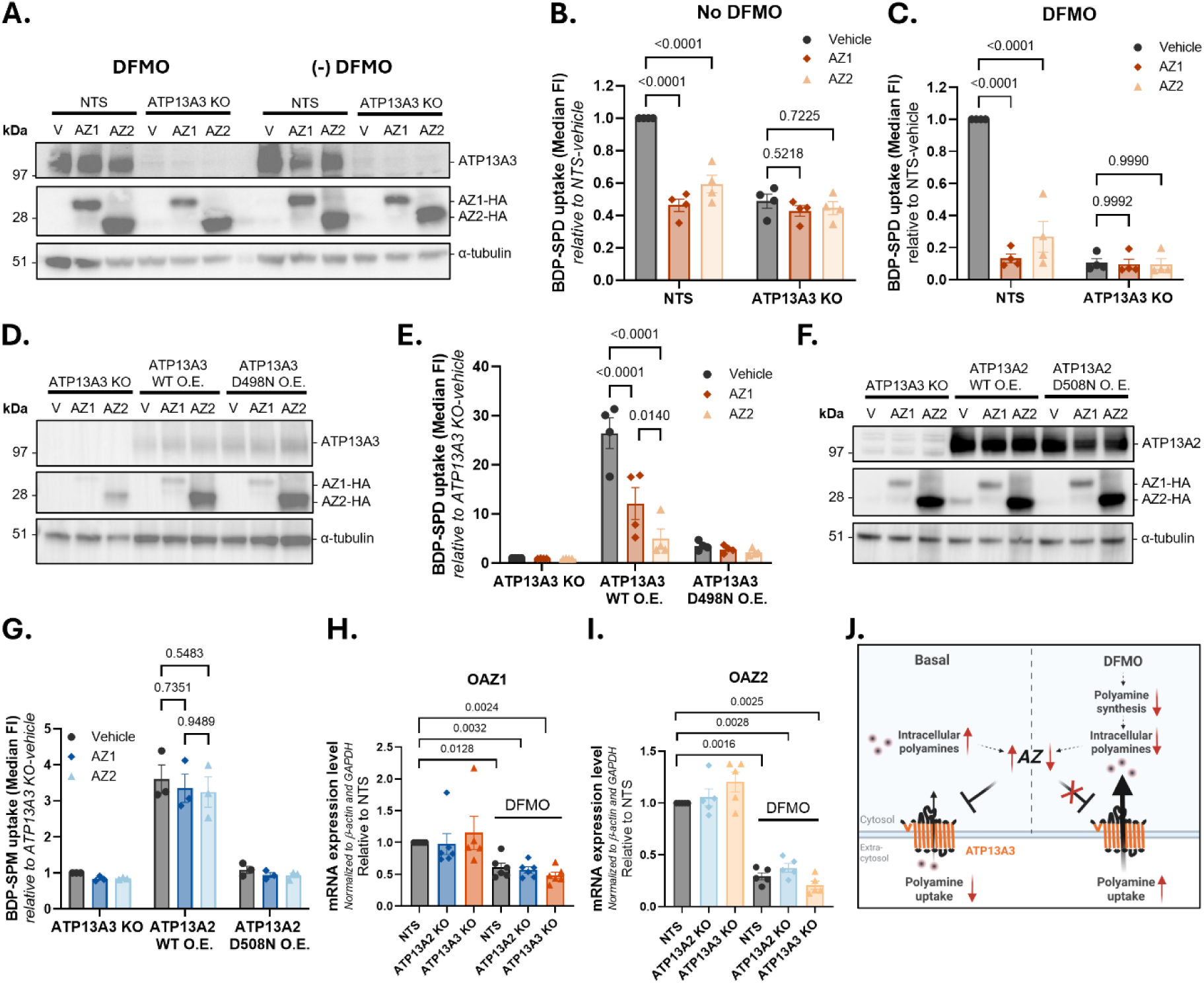
Antizyme selectively inhibits ATP13A3-mediated polyamine uptake in cells. **(A-C)** Effect of antizyme (AZ) expression on ATP13A3-mediated polyamine uptake. **(A)** Representative immunoblots showing ATP13A3, AZ1-HA, and AZ2-HA expression in HEK293T NTS and ATP13A3 knockout (KO) cells transfected with Fugene HD (V), AZ1, or AZ2 in the absence or presence of 20 h pre-treatment with 1.5 mM DFMO. α-Tubulin served as a loading control. **(B-C)** BODIPY-spermidine (BDP-SPD; 0.1 µM, 30 min) uptake in the corresponding cell lines and conditions shown in **(A)**, in the absence **(B)** or presence **(C)** of pre-treatment with 1.5 mM DFMO for 20 h. Data are presented as mean ± SEM of *n* = 4 independent biological replicates, with individual data points shown. Two-way ANOVA with Tukey’s multiple comparisons test. **(D-E)** Effect of AZ expression on ATP13A3-mediated polyamine uptake in ATP13A3 rescue cell lines. **(D)** Representative immunoblots showing ATP13A3, AZ1-HA, and AZ2-HA expression in ATP13A3 KO cells and ATP13A3 KO cells stably re-expressing ATP13A3 WT or the catalytically inactive mutant D498N, transfected with Fugene HD (V), AZ1, or AZ2 and pre-treated with 1.5 mM DFMO for 20 h. α-Tubulin served as a loading control. **(E)** BDP-SPD (1 µM, 30 min) uptake in the corresponding cell lines and conditions shown in **(D)**. Data are presented as mean ± SEM of *n* = 4 independent biological replicates, with individual data points shown. Two-way ANOVA with Tukey’s multiple comparisons test. **(F-G)** Effect of AZ expression on ATP13A2-mediated polyamine uptake. **(F)** Representative immunoblots showing ATP13A2, AZ1-HA, and AZ2-HA expression in ATP13A3 KO cells and ATP13A3 KO cells stably re-expressing ATP13A2 WT or the catalytically inactive mutant D508N, transfected with Fugene HD (V), AZ1, or AZ2 and pre-treated with 1.5 mM DFMO for 20 h. α-Tubulin served as a loading control. **(G)** BDP-spermine (BDP-SPM; 1 µM, 30 min) uptake in the corresponding cell lines and conditions shown in **(F)**. Data are presented as mean ± SEM of *n* = 3 independent biological replicates, with individual data points shown. Two-way ANOVA with Tukey’s multiple comparisons test. **(H-I)** *OAZ1* **(H)** and *OAZ2* **(I)** mRNA expression in HEK293T NTS, ATP13A2 KO and ATP13A3 KO cells in the presence of absence of pre-treatment with 1.5 mM DFMO for 24 h. Data are presented as relative expression (2^−ΔΔCt). Ct values were normalized to β-actin (ACTB) and GAPDH, and expression was normalized to NTS. Bars represent mean ± SEM of *n* = 4-5 independent biological replicates, with individual data points shown. Statistical analyses were performed on ΔCt values using one-way ANOVA followed by Dunnett’s multiple comparisons test. **(J)** Proposed model of antizyme-dependent regulation of ATP13A3-mediated polyamine uptake. Under basal conditions, elevated intracellular polyamine levels induce antizyme (AZ), which inhibits ATP13A3-mediated polyamine uptake. Following DFMO treatment, inhibition of polyamine biosynthesis reduces intracellular polyamine levels, relieving AZ-dependent inhibition and promoting ATP13A3-mediated polyamine uptake to restore polyamine homeostasis. Created with Biorender.

We next used a reconstitution strategy in HEK293T cells with ATP13A3 KO to further determine whether antizyme impacts ATP13A3 specifically (**Fig. 4D-E**). Re-expression of WT ATP13A3 restored antizyme-sensitive polyamine uptake, whereas ATP13A3-D498N did not, pointing to an ATP13A3 transport-dependent antizyme regulation (**Fig. 4E**). AZ1 and AZ2 reduced ATP13A3-mediated BODIPY-spermidine uptake by approximately 2.3- and 4.7-fold (**Fig. 4E**), respectively, but failed to suppress uptake in ATP13A3 KO cells expressing ATP13A2 WT or ATP13A2-D508N (**Fig. 4F-G**), which confirms the ATP13A3 selective regulation by antizyme in a cellular context. Note that AZ1 was expressed at lower levels than AZ2 (**Fig. S9C**), but both isoforms inhibited uptake of ATP13A3, and not ATP13A2, supporting a potent regulatory effect of antizyme on ATP13A3-mediated transport. Together, our cellular polyamine uptake experiments place ATP13A3 downstream of antizyme in the DFMO-induced uptake response. When polyamine synthesis was blocked in HEK293T cells by DFMO, antizyme expression was significantly reduced (**Fig. 4H-I**), together with a modest, but non-significant increase in AZIN2 expression, whereas AZIN1 expression remained unaffected (**Fig. S9D-E**). The negative impact of DFMO on antizyme levels therefore provides a mechanistic explanation of how polyamine uptake is stimulated via ATP13A3. Indeed, forcing antizyme expression under DFMO conditions reinstates the inhibitory brake on ATP13A3-mediated polyamine uptake even under polyamine-depleted conditions in HEK293T cells (**Fig. 4C**).

In conclusion, antizyme selectively inhibits cellular polyamine uptake via ATP13A3, which is lifted under DFMO (**Fig. 4J**). The antizyme-ATP13A3 axis therefore represents the regulatory switch controlling adaptive polyamine uptake under DFMO treatment.

### AMXT 1501 pharmacologically blocks ATP13A3 via a competitive mechanism

The identification of ATP13A3 as the dominant DFMO-responsive polyamine importer and as direct binding partner of antizyme suggests that this transporter could be pharmacologically targeted for polyamine depletion therapy in cancer. AMXT 1501 (**Fig. 5A**) was developed to block compensatory polyamine uptake during DFMO-based polyamine depletion therapy, but its target and mode of action has remained incompletely established (*16*).

**Figure 5:**
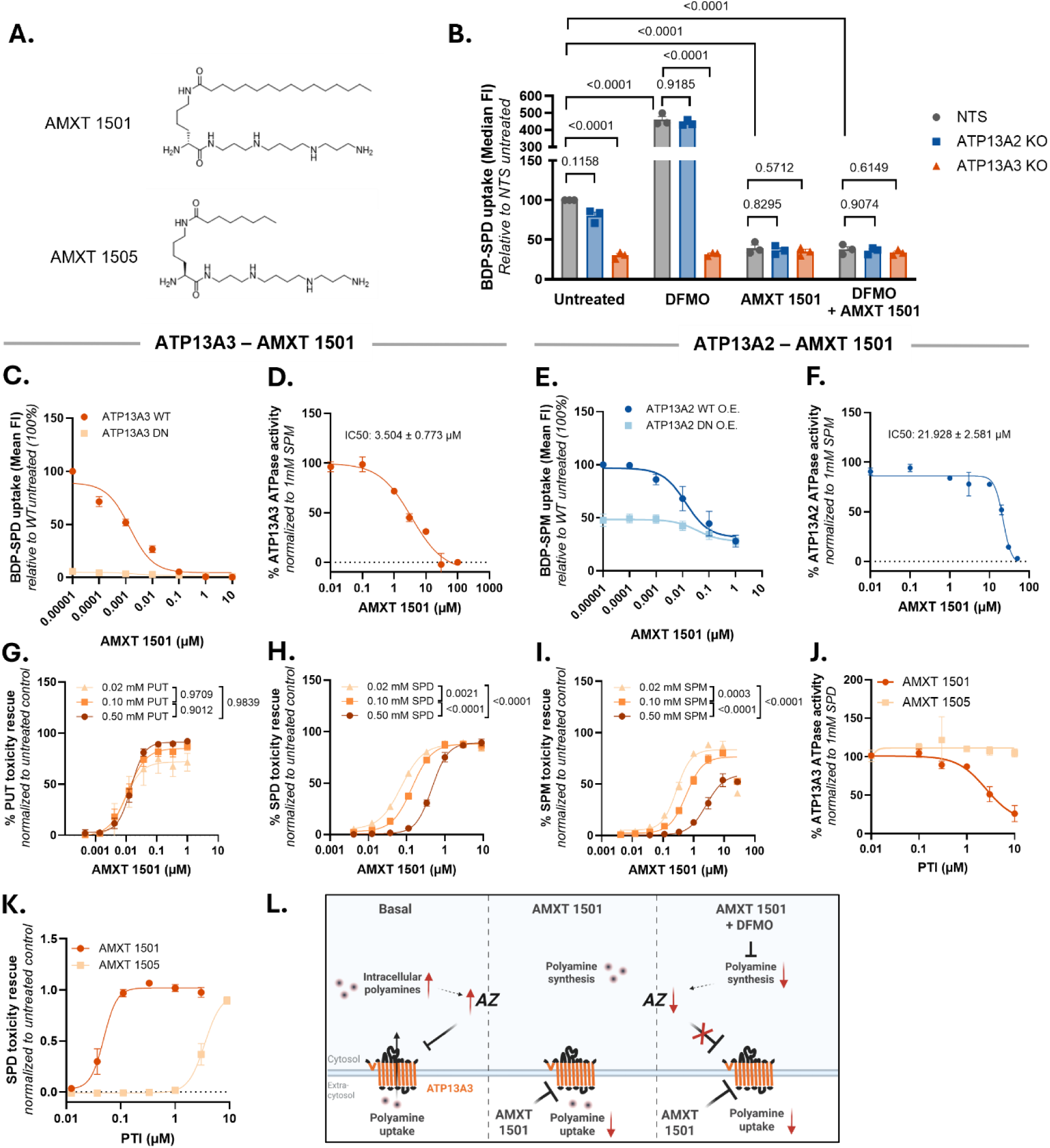
AMXT 1501 inhibits ATP13A3 activity under basal and DFMO conditions. **(A)** Molecular structure of AMXT 1501 and a shorter lipid tail analog AMXT 1505. **(B)** BODIPY-spermidine (BDP-SPD; 0.1 µM, 30 min) uptake in HEK293T NTS cells, ATP13A2 KO and ATP13A3 KO cells. Cells were pretreated for 24h with or without 1.5 mM DFMO and/or for 2h with or without 1 µM AMXT 1501. Uptake was quantified as median fluorescence intensity and normalized to untreated NTS HEK293T cells, set to 100%. Data are presented as mean ± SEM, with individual data points shown. *n* = 3 independent experiments. Statistical analyses were performed on log₁₀-transformed raw uptake values using two-way ANOVA followed by Tukey’s multiple-comparisons test. Normality was assessed using a QQ plot. **(C)** Uptake of 1 µM BODIPY-conjugated spermidine (BDP-SPD; 1 µM 30 min, 1 mM aminoguanidine) in SH-SY5Y cells with wild type ATP13A3 overexpression (ATP13A3 WT O.E.) or overexpression of the catalytically dead D498N mutant (ATP13A3 D498N O.E.). Uptake was quantified as mean fluorescence intensity and normalized to untreated ATP13A3 WT O.E. SH-SY5Y cells, set to 100%. Data are presented as mean ± SEM of *n* = 3 independent biological replicates, with individual data points shown. **(D,F)** Spermine (SPM)-stimulated ATPase activity of purified ATP13A3 **(D)** or ATP13A2 **(F)** in the presence of increasing concentrations of AMXT 1501. ATPase activity was normalized to the activity measured at 1 mM SPM in the absence of AMXT 1501. Data are presented as mean ± SEM of *n* = 3 independent biological replicates. **(E)** Uptake of BODIPY-conjugated spermine (BDP-SPM; 1 µM 30 min) in SH-SY5Y cells with wild type ATP13A2 overexpression (ATP13A2 WT O.E.) or overexpression of the catalytically dead D508N mutant (ATP13A2 D508N O.E.), following the same experiment procedure described in **(C)**. **(G-I)** Competition experiments between AMXT 1501 and polyamines putrescine (PUT) **(G)**, spermidine (SPD) **(H)**, or SPM **(I)** in SH-SY5Y cells overexpressing WT ATP13A3. Excessive extracellular polyamine uptake in SH-SY5Y cells overexpressing WT ATP13A3 leads to toxicity by intracellular polyamine overload which can be blocked with AMXT 1501. Cells were pre-treated for 1h with increasing concentrations of AMXT 1501 before incubating with toxic polyamine concentrations overnight to induce toxicity. Cell viability was measured using the MUH-fluorescence-based cell viability assay and quantified by background subtraction and normalization to untreated cells. Data are presented as mean ± SEM of *n* = 5 independent biological replicates, with individual data points shown. Log-transformed IC₅₀ values were compared among groups using two-way ANOVA followed by Tukey’s multiple-comparison test. **(J)** SPD-stimulated ATPase activity of purified ATP13A3 in the presence of increasing concentrations of AMXT 1501 or AMXT 1505. ATPase activity was normalized to the activity measured at 1 mM SPD in the absence of AMXT 1501. Data are presented as mean ± SEM of n = 3 independent biological replicates. **(K)** SPD-induced toxicity rescue in HEK293T ATP13A3 KO cells overexpressing ATP13A3 with AMXT 1501 and shorter lipid tail analog AMXT 1505. HEK cells were pretreated for 2h with increasing concentrations of Polyamine transport inhibitors (PTIs) after which toxicity was induced with 0.1 mM SPD. After overnight incubation, cell viability was measured using using the CellTiter-Glo® Luminescent Cell Viability Assay. Data were prepared for plotting by subtracting the background luminescence and normalizing to untreated cells. Data are presented as mean ± SEM, with individual data points shown. n = 4. **(L)** Proposed model of inhibition of ATP13A3-mediated polyamine uptake by AMXT 1501. Under basal conditions, elevated intracellular polyamine levels induce expression of antizyme (AZ), which blocks ATP13A3-mediated polyamine uptake. The polyamine transport inhibitor AMXT 1501 directly inhibits ATP13A3-mediated polyamine uptake under both basal and α-difluoromethylornithine (DFMO)-treated conditions. Created with Biorender.

In HEK293T cells, AMXT 1501 had no functional impact in the absence of ATP13A3, whereas AMXT 1501 lowered uptake in ATP13A2 KO cells to the level of ATP13A3 KO in basal and DFMO conditions (**Fig. 5B**), consistent with inhibition of the prevailing polyamine uptake mechanism under ATP13A3 control (**Fig. 1**). We further demonstrated that AMXT 1501 inhibited ATP13A3-mediated polyamine uptake in SH-SY5Y cells (**Fig. 5C**) as well as the polyamine-dependent ATPase turnover of purified ATP13A3 (**Fig. 5D, Fig. S10A**). This inhibition was not selective for ATP13A3, since AMXT 1501 also inhibited ATP13A2-dependent polyamine uptake (**Fig. 5E**) and ATPase activity (**Fig. 5F**) albeit with lower potency, indicating that AMXT 1501 is a broad-specificity polyamine transport inhibitor. In an orthogonal assay, AMXT 1501 rescued the toxicity in ATP13A3-overexpressing cells exposed to excessive extracellular polyamine concentrations, which cause toxic intracellular polyamine overload (**Fig. 5G-I**). We further demonstrated that the protective effect of AMXT 1501 decreased with higher extracellular spermidine or spermine concentrations (**Fig. 5H-I**), suggesting a competitive inhibition mechanism, consistent with competition at the conserved polyamine-binding cavity of ATP13A2 and ATP13A3 (**Fig. S4B-C**). AMXT 1505, a variant of AMXT 1501 with a shorter lipophilic tail (**Fig. 5A**) was markedly less effective in the inhibition of ATP13A3 ATPase activity (**Fig. 5J, Fig. S10B**) and in protecting cells from spermidine toxicity (**Fig. 5K**).

Thus, AMXT 1501 pharmacologically inhibits ATP13A3 via a direct competitive inhibition mechanism, thereby preventing the main polyamine uptake in cells under ATP13A3 control, both in basal and DFMO conditions (**Fig. 5L**).

### The ATP13A3-antizyme pathway is promoted in multiple cancer types and progression states

Since AMXT 1501 is currently explored in cancer clinical trials in combination with DFMO (NCT07287917, NCT06465199) (*15*), we next examined the pathological relevance of the antizyme-ATP13A3 axis in cancer disease. Elevated ATP13A3 expression has previously been associated with worse prognosis in neuroblastoma (*16*), breast cancer (*31*), pancreatic cancer (*23, 38*), and head and neck cancer (*30*), whereas ATP13A2 expression may be linked to a better outcome (*16, 31*). With the new mechanistic insights on ATP13A3-antizyme control, we have revisited publicly available patient transcriptomics datasets of 12 cancer types to assess the housekeeping genes involved in polyamine acquisition and its regulation (*i.e. ATP13A2/3*, *ODC1*, *OAZ1/2*, *AZIN1*). All 12 investigated cancer types revealed gene-expression signatures consistent with increased polyamine acquisition across cancers, which therefore emerges as a more general cancer strategy. However, the reprogramming of polyamine homeostasis differed between cancer types (**Fig. 6A, Fig. S11**) and progression states (**Fig. 6B, Fig. S12A-B**). Some cancer types relied more on *ODC1* upregulation (3/12 cancer types), others were marked by upregulated *ATP13A3* expression (3/12), but the majority of cancer types (6/12) belonged to a third group where both arms were simultaneously promoted (**Fig. 6A**). Significant positive correlations between *ATP13A3*/*AZIN1* (in 12/12 cancer types) and *ATP13A3/ODC1* (11/12) were observed, whereas a negative correlation was found for *ATP13A3*/*OAZ1* (10/12), but not *ODC1*/*OAZ1* (**Fig. S11**), pointing to a dysregulated polyamine uptake control in all cancer types. Although melanoma belonged to the cancer types with elevated *ODC1* (**Fig. 6A**; SKCM) we observed dynamic changes of metastatic melanoma progressing more towards a polyamine uptake fingerprint in two independent datasets (**Fig. 6B, Fig. S12A**). Indeed, *ATP13A3* and *AZIN1* mRNA expression, were significantly higher in both datasets of metastatic melanoma (**Fig. 6B, Fig. S12A**). *ODC1* mRNA was significantly higher, whereas *OAZ1* and *OAZ2* mRNA were lower in only one of the two datasets (**Fig. S12A)**. Further signs of plasticity towards a more prominent polyamine uptake profile were observed during melanoma drug exposure and resistance. We observed lower *ODC1*, equal *ATP13A3*, but lower *OAZ1* mRNA levels, which predict a stronger reliance on polyamine uptake (**Fig. S12B**).

**Figure 6:**
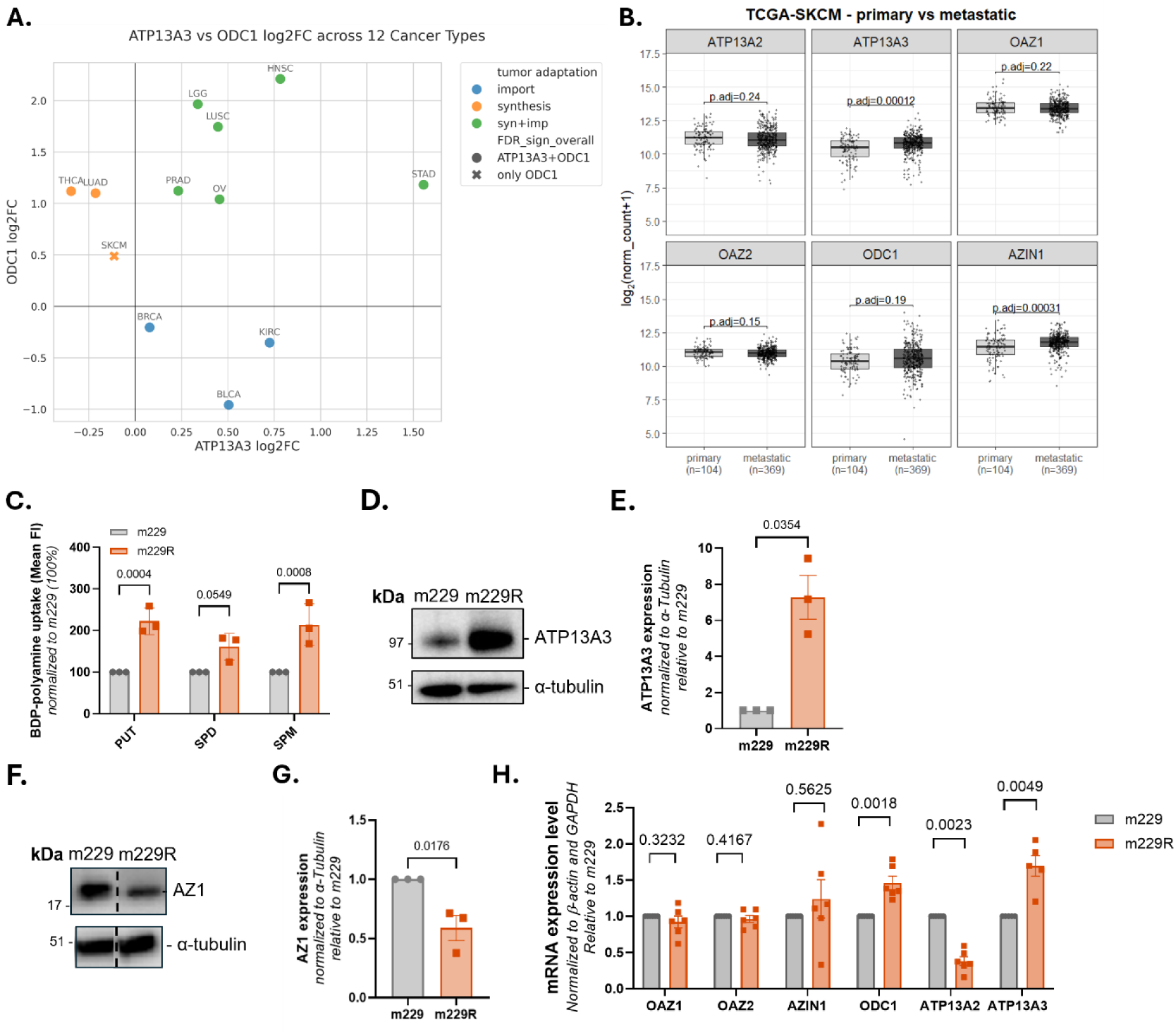
Polyamine acquisition is promoted across cancer types and melanoma progression. **(A)** Log2 fold changes in mRNA expression of *ATP13A3* and *ODC1* are plotted for 12 common cancer types (relative to their matched normal tissues). Log2 fold changes were calculated as the difference in arithmetic means between tumor and normal samples in log2 space. Tumor adaptation (synthesis + import; synthesis only; import only) is color-coded. Statistical significance is coded by marker style. All log2FC values were statistically significant (BH-FDR) to p <0.001 except for *ATP13A3* in BRCA (p < 0.05) and *ATP13A3* in SKCM (not significant). **(B)** The Cancer Genome Atlas (TCGA) Skin Cutaneous Melanoma (SKCM) was used to compare mRNA expression [UCSC, Xena, log₂(normalized count + 1)] of *ATP13A2*, *ATP13A3*, *OAZ1-2*, *ODC1* and *AZIN1* between primary (n=104) and metastatic (n=369) tumours. Groups were compared by Welch t-test with Benjamini-Hochberg correction (adjusted p-values shown). **(C)** Uptake of 1 µM BODIPY-putrescine (BDP-PUT), BODIPY-spermidine (BDP-SPD), and BODIPY-spermine (BDP-SPM), in melanoma m229 or m229 resistant (m229R) cells. Cells were treated for 30 min with BODIPY-polyamines in the presence of 1 mM aminoguanidine. Uptake was quantified as mean fluorescence intensity and normalized to m229 cells. Data are presented as mean ± SEM of n= 3 independent biological replicates, with individual data points shown. Representative immunoblot **(D)** and quantification **(E)** showing ATP13A3 expression in m229 and m229R cells. α-Tubulin served as a loading control. Data are presented as mean ± SEM of n = 3 independent biological replicates. Unpaired t test with Welch’s correction. Representative immunoblot **(F)** and quantification **(G)** showing antizyme 1 (AZ1) expression in m229 and m229R cells α-Tubulin served as a loading control. The blot was cropped for presentation, with the dividing line marking the position of the removed lanes. Data are presented as mean ± SEM of n = 3 independent biological replicates. Unpaired t test. **(H)** mRNA expression of antizymes *OAZ1* and *OAZ2*, antizyme inhibitor 1 (*AZIN1*), *ODC1*, *ATP13A2* and *ATP13A3* in m229 and m229R cells. Data are presented as relative expression (2^−ΔΔCt). Ct values were normalized to *ACTB* and *GAPDH*, and expression was normalized to m229. Bars represent mean ± SEM of *n* = 6 independent biological replicates, with individual data points shown. Statistical analyses were performed on ΔCt values using a paired two-tailed t-test or the Wilcoxon signed-rank test, after assessing normality using a QQ plot.

Since mRNA expression only gives a partial answer, we experimentally confirmed that the antizyme-ATP13A3 axis was promoted in two common *in vitro* melanoma models of tumor plasticity in response to vemurafenib therapy (a BRAF inhibitor), *i.e.* the human melanoma cell pairs m229P/R and m238P/R (P, parental = non-resistant to vemurafenib; R, resistant to vemurafenib) (**Fig. 6C, Fig. S12C**). Both m229R (**Fig. 6C**) and m238R (**Fig. S12C**) exhibited higher polyamine uptake as compared to their respective parental cell line. In m229R, we demonstrated that this coincides with increased ATP13A3 protein levels (**Fig. 6D-E**) and reduced AZ1 protein levels (**Fig. 6F-G**). At the mRNA level, we did not observe *OAZ* changes, but observed reduced *ATP13A2* expression and significantly elevated *ATP13A3* and *ODC1* expression (**Fig. 6H**).

In conclusion, the distinct polyamine acquisition fingerprints among cancer types and progression states illustrate that cancer cells exploit various complementary mechanisms to reprogram polyamine homeostasis towards a higher polyamine load, of which ATP13A3-antizyme regulation is a newly identified pathway. The dynamic changes between polyamine uptake and synthesis pathways provide a strong rationale for the ongoing clinical trials with AMXT 1501 (*15*) and DFMO combination therapy in melanoma and other cancer types.

## Discussion

### Antizyme regulates polyamine homeostasis by coupling polyamine sensing to coordinated polyamine acquisition

As essential nitrogen-rich metabolites that support growth but become toxic in excess, polyamines require stringent control of their intracellular availability via synthesis and uptake (*2*). Here, we resolve a critical missing link of the polyamine feedback circuit by demonstrating that antizyme inhibits cellular polyamine uptake through ATP13A3. Polyamine synthesis and uptake are not autonomous pathways, but represent coordinated outputs of a common negative-feedback circuit via antizyme (*5, 8*). The *OAZ* to AZ translational circuit couples the free cytosolic polyamine pool to antizyme protein expression. A rise in cytosolic polyamine concentration increases full-length antizyme protein via an *OAZ* translational circuit that relies on +1 ribosomal frameshifting of *OAZ* mRNAs (*39, 40*) and via polyamine binding to nascent antizyme, which promotes completion of translation (*41*). Once synthesized, antizyme suppresses ODC1, the rate-limiting step in polyamine synthesis, by disrupting the functional ODC1 homodimer leading to ODC1 inactivation and degradation (*7*).

We here demonstrate that AZ1 and AZ2 selectively inhibit ATP13A3, a bona fide polyamine transporter that we establish as a major contributor to cellular polyamine uptake, while sparing ATP13A2. Although both transporters exhibit similar properties and contribute to intracellular polyamine availability and distribution, they control separate polyamine fluxes that are differentially regulated. The selective antizyme inhibition of ATP13A3 allows for separate control of ATP13A3-mediated polyamine import at the plasma membrane and early/recycling endosomes (*23, 42*) *versus* the ATP13A2-mediated export of polyamines out of the late endo-lysosomes (*20*). The latter may represent a homeostatic mechanism for local polyamine delivery (*20, 27, 43, 44*) and/or polyamine recycling during autophagy (*45*). Conversely, their overlapping properties do not exclude that they may act in complementary or partially redundant ways, which may explain why ATP13A3 can compensate for ATP13A2 loss in disease models of ATP13A2 dysfunction when spermidine is supplied (*43*). Hence, ATP13A3 may provide a compensatory role in ATP13A2-associated parkinsonism, which fits with their overlapping biochemical properties. The differential regulation of ATP13A2 and ATP13A3 together with their distinct subcellular localization and impact on polyamine homeostasis provides a rationale for their distinct (patho)physiological roles (*3*).

Based on our results and other published findings (*46*), ATP13A3 and ODC1 interact with AZ1/2 via an overlapping, conserved AZ1/2 surface that is also the binding site for AZ-I1 (*46*), suggesting that not only ATP13A3 and ODC1, but also AZ-I1 may compete for antizyme binding to provide a coordinated control of polyamine synthesis and uptake. Indeed, AZ-I1/2 isoforms positively modulate the uptake of extracellular polyamines (*47, 48*), which fits well with the established ATP13A3-antizyme interface that is also used for interaction with ODC1, and hence also for AZ-I1 and AZ-I2 (*7*). Such a multilevel control system via antizymes and AZ-I1/2 may function as a tunable rheostat, rather than a simple on/off switch, allowing both homeostatic stability and flexibility to reconfigure polyamine acquisition quickly to adapt polyamine availability to demand.

ATP13A3-antizyme control explains how cells coordinate polyamine sensing and exogenous polyamine supply, which has broad implications to understand polyamine homeostasis and its alterations during disease and therapeutic intervention. The balance between synthesis, uptake, and recycling is likely species-, tissue- and context-dependent. A change in balance may be achieved by altering the relative expression of the antizyme isoforms AZ1-2 and their binding partners ODC1, ATP13A3, and AZ-I1/2, but may also depend on their spatial organization and relative binding affinities for antizyme (*49*).

In an *in vivo* context (*Caenorhabditis elegans*), polyamine uptake contributes more to polyamine levels than polyamine synthesis (*50*). Spermidine supplementation offers cardiovascular (*51*) and neurological protection (*52*), countering the decline in polyamine levels that occurs with ageing. Although the mechanisms behind this decline have not been resolved, a disturbed ATP13A3-antizyme regulation may play a role. Polyamine uptake pathways are further important for polyamine homeostasis, since genetic loss of *ATP13A3* has been genetically associated with pulmonary arterial hypertension, a rare cardiovascular disorder (*24, 29*), whereas an excessive ATP13A3 activity may play a role in cancer.

### ATP13A3-antizyme control is disturbed in multiple cancer types and progression states

In this study, we demonstrated that increased ATP13A3 expression and/or its regulation of activity is implicated in various cancers and cancer progression. Proliferating cells are highly dependent on polyamines to provide sufficient charge-neutralizing polyamines for DNA and RNA stability, chromatin reorganization, ribosome assembly, as well as for hypusination of eIF5A, a polyamine-dependent post-translational modification required for growth-associated translation. Cells therefore increase polyamine supply through ODC1-driven *de novo* synthesis, often induced by growth-factor and MYC/MYCN programs, (*4, 53*) and/or by stimulating uptake of extracellular polyamines, which is mainly ATP13A3 reliant.

The ATP13A3-antizyme control provides a newly identified mechanism that allows cancer cells to upregulate polyamine acquisition, which goes beyond already established mechanisms involving AZ-I1/2, AZ1/2, and ODC1 (*4–6*). Importantly, all cancer types show significant reprogramming of the polyamine homeostasis towards elevated polyamine acquisition, which therefore emerges as a more general, recurrent strategy for sustaining polyamine availability in cancer. However, the underlying mechanisms differ between cancer types and progression state. Indeed, we revealed plasticity of the polyamine acquisition system during melanoma progression and/or drug resistance, suggesting that tumor evolution can dynamically rewire polyamine supply, moving between synthesis-dominant, uptake-dominant, and dual synthesis–uptake states. This is further illustrated by the DFMO-induced increase in polyamine uptake, which we here have mechanistically explained by the feedback system of antizyme on ATP13A3. DFMO-induced uptake is the predictable consequence of releasing ATP13A3 from antizyme-mediated feedback control.

The dynamic reprogramming of polyamine acquisition in cancer cells explains why targeting only the polyamine synthesis has had only limited therapeutic success. By placing ATP13A3 downstream of antizyme and most likely AZ-I1 control, we established ATP13A3 as an interesting drug target to block polyamine uptake. To prevent dynamic reprogramming of polyamine homeostasis during treatment, a dual polyamine-depletion approach by combining ODC1 and ATP13A3 inhibitors will be most efficient. This strategy inhibits the two main polyamine acquisition pathways, the rate limiting step ODC1 in polyamine synthesis and the main polyamine importer ATP13A3.

Note that our cancer-dataset analyses are based primarily on mRNA abundance and therefore identify potential pathway states rather than directly measuring protein levels or polyamine-uptake. This distinction is particularly important for the polyamine pathway, in which ODC1, antizymes, and AZINs are extensively regulated at translational and post-translational levels, of which several mechanisms fall under oncogene control. Nevertheless, a large body of work has previously established that ODC1 and the AZ/AZ-I1 regulation module are disturbed across different cancer types, strongly supporting the significance of the altered mRNA expression levels observed throughout this pathway (*4–6*). By defining ATP13A3 as an antizyme-regulated polyamine uptake transporter, our study extends this framework from polyamine synthesis to the broader concept of polyamine acquisition, also involving polyamine uptake. However, how ATP13A3 transcript and/or protein expression may be regulated remains to be established. Analysis of ATP13A3 transporter expression may be considered as a biomarker (*30, 31, 38*), but our work implies that a combination with other genes involved in polyamine accumulation would provide a more comprehensive view of how cancer cells acquire polyamines. Importantly, in melanoma models, we have validated this concept beyond transcript analysis by showing concordant alterations at the protein and functional levels in parental *versus* drug resistant melanoma cell models. This illustrates that dataset-derived signatures may reflect biologically meaningful changes in polyamine homeostasis.

### Implications for cancer therapy

Our mechanistic insights on polyamine uptake control and pharmacological intervention are timely and provide the rationale for ongoing drug development efforts towards polyamine depletion strategies. DFMO has recently been clinically approved to reduce relapse risk of neuroblastoma (*11*). Conversely, several polyamine transport inhibitors have been developed based on their capacity to block cellular polyamine uptake (*13, 14, 18, 54*), although a clear definition of the polyamine transporter(s) that are targeted has slowed down further on target development. By conclusively establishing ATP13A3 as a main target of AMXT 1501, we here resolve a mode of action mechanism for this compound class and provide target validation for ATP13A3 in polyamine depletion strategies. Our study provides a strong rationale for clinical testing of AMXT 1501 and DFMO combination therapy, which is currently under investigation for advanced solid tumors, metastatic melanoma, ER-positive/HER2-negative breast cancer, pediatric neuroblastoma, CNS tumors, and sarcomas NCT07287917, NCT06465199, NCT05717153).

Therapeutic development of ATP13A3 antagonists will also require careful attention to selectivity and safety. ATP13A3 has physiological roles outside cancer, and rare loss-of-function ATP13A3 variants have been genetically associated with pulmonary arterial hypertension (*24, 29, 55*), indicating that sustained loss of ATP13A3 function may have tissue-specific consequences, although environmental triggers may also play a role (*56*). Our study also establishes AMXT 1501 as a broad-specificity, competitive inhibitor of both ATP13A3 and ATP13A2, although in cells, its effect on polyamine uptake may mostly depend on ATP13A3 inhibition. Initial Phase I results of AMXT 1501/DFMO combination therapy suggest safety and tolerability with evidence of preliminary clinical activity (*15*). This raises the question to which extent ATP13A3 selectivity is needed for antitumor activity and how much ATP13A2 inhibition can be tolerated. These questions are especially relevant for brain-penetrant inhibitors that could be valuable for glioma and other brain tumors, but it may also require monitoring for ATP13A2-related liabilities. Whether other polyamine transport inhibitors under investigation work on ATP13A3 and/or present differences in selectivity remains unknown (*19*). By firmly establishing ATP13A3 as the most relevant drug target for blocking polyamine uptake, our studies open the door for the development of more ATP13A3-selective and/or allosteric inhibitors.

Together, our findings define ATP13A3 as the main antizyme-regulated polyamine uptake transporter and a pharmacological target of AMXT 1501. This completes a major missing branch of the polyamine homeostatic circuit that is exploited by cancer cells to increase polyamine supply. Dysregulation of the ATP13A3-antizyme feedback control creates a common and targetable vulnerability in cancer. Our study provides a molecular framework for understanding DFMO resistance, for monitoring polyamine dyshomeostasis in patients, developing next-generation polyamine-transport inhibitors, and for testing dual polyamine-depletion strategies in metastatic and therapy-resistant cancers.

## Resource and materials availability

Further information and requests for resources and reagents should be directed to and will be fulfilled by the corresponding author, Peter Vangheluwe. This study did not generate new unique reagents. All experimental protocols are available via protocols.io. Details of protocols, datasets, software, reagents, and cell lines are provided in the Key Resource Table.

## Author Contributions

The study was supervised and coordinated by P.V. with A.Sc., J.C., R.A., J.E. and S.v.V. contributing to the design and providing critical input; P.A. provided input and resources for the melanoma research; St.Ve. provided input and resources for the AMXT 1501 work; A.Sc., R.A. and J.B. executed the antizyme study; J.C., S.v.V., L.D., M.D. and E.A. performed the ATP13A2/3 biochemical characterization; E.R. and A.Sc. validated and characterized cell models; A.Sc., J.C. and L.D. executed the AMXT 1501 work; St.Vr., E.R. and M.A. performed radiolabeled uptake experiments; Y.F., E.R. and E.M. executed the melanoma experimental work; J.B., A.V., L.D., A.Sh. E.A., N.L. and M.D. provided general support and performed experiments; S.C. conducted the lipid binding assays; J.E. and E.M. performed in silico analyses of human expression datasets; C.V.d.H. generated stable cell lines; C.V.d.H. and R.G. developed the CRISPR/Cas9 virus-like particle strategy; N.M. provided protocols and constructs for antizyme 2 purification; P.V. secured funding; S.v.V., A.Sc., M.A. compiled the first draft of the manuscript, which was finalized by P.V., and reviewed, edited and approved by all authors.

## Data Availability

All data, protocols and lab material are listed on the Key Resource Table with the respective DOI or identifiers at DOI: 10.5281/zenodo.21102857. All data generated or analysed in this study can be found through the Zenodo repository at DOI: 10.5281/zenodo.21102857. All protocols are shared via protocols.io at (Bring structure to your research - protocols.io). All data and protocols will be made publicly available as of the date of publication.

## Code availability

All original code has been deposited at GitLab (https://gitlab.kuleuven.be/lcts/lcts_papers) and is publicly available. The macro for Fig. 6 A-B, Fig. S11, Fig. S12 A-B has been deposited in the Zenodo database under accession code 10.5281/zenodo.21099337 (https://doi.org/10.5281/zenodo.21099337).

## Open Access

This article is licensed under a Creative Commons Attribution 4.0 International License, which permits use, sharing, adaptation, distribution and reproduction in any medium or format, as long as you give appropriate credit to the original author(s) and the source, provide a link to the Creative Commons licence, and indicate if changes were made. The images or other third party material in this article are included in the article’s Creative Commons licence, unless indicated otherwise in a credit line to the material. If material is not included in the article’s Creative Commons licence and your intended use is not permitted by statutory regulation or exceeds the permitted use, you will need to obtain permission directly from the copyright holder. To view a copy of this licence, visit http://creativecommons.org/licenses/by/4.0/.

## Acknowledgments

Cartoon elements of figure panels were created using BioRender.com. We acknowledge our frequent use of the facilities and equipment of the Leuven Viral Vector Core facility (KU Leuven) and the KU Leuven FACS Core. We thank Mark Burns for the generous supply of AMXT 1501 and AMXT 1505.

## Materials and methods

### Cell lines and culture

SH-SY5Y cells (CRL-2266, RRID:CVCL_0019) and HEK293T cells (CRL-11268, RRID:CVCL_1926) were purchased from ATCC. The human melanoma M229P/M229R and M238P/M238R isogenic cell lines, were a generous gift from R. Lo (UCLA). M238P/M238R isogenic cell lines were a generous gift from R. Lo (UCLA). M229 and M238 cells were cultured in DMEM supplemented with 10% heat-inactivated fetal bovine serum (Pan Biotech), 1% sodium pyruvate (Gibco) and 1% penicillin-streptomycin (Sigma). The M229 and M238 resistant cells (M229R/M238R) were cultured in the same medium with 1µM PLX4032 (Vemurafinib, Cayman) to maintain resistance as described in (*57*).

SH-SY5Y cells were cultured in high glucose Dulbecco’s modified Eagle’s medium (DMEM, Gibco) supplemented with 15% heat-inactivated FBS (Pan Biotech), 1% non-essential amino acids (Sigma), 1% sodium pyruvate (Sigma) and 2% penicillin-streptomycin (Sigma). HEK293T cells were cultured in DMEM supplemented with 8% heat-inactivated FBS (Pan Biotech) and 2% penicillin-streptomycin (Sigma). All cell lines described above were maintained at 37 °C in a humidified incubator with 5% CO₂. Routine mycoplasma testing was performed. Heat-inactivated FBS was used to deplete polyamine oxidase activity, which could interfere with fluorescent polyamine probes or supplemented polyamines.

ATP13A2 and ATP13A3 KO cell line generation – HEK293T ATP13A2 and ATP13A3 knockout (KO) cell lines were generated using virus-like particles (VLPs) as previously described(*58*). The following VLPs and their corresponding sequences were used: sgRNA1 targeting hATP13A3 exon 4 with guide sequence: ACTCAGGCATCCAATAGAGG (used in antizyme experiments), sgRNA3 targeting hATP13A3 exon 16 with guide sequence: AATGTTTGAGGCTATTGGAT (used in characterization of uptake and AMXT 1501 treatments), sgRNA targeting hATP13A2 with guide sequence: CGTCAGGGTCCCATAACCGG, and empty VLP without guide but only cas9. KO was confirmed by sequencing of genomic DNA, qPCR and western blotting. Clones with confirmed protein KO were pooled.

### Stable cell line generation

Stable cell lines overexpressing human ATP13A2 isoform 2 (NM_001141973.3; wild-type, Addgene plasmid #171485, or the catalytically inactive D508N mutant, Addgene plasmid #171820) or human ATP13A3 isoform 1 (NM_024524.4; wild-type Addgene plasmid #259310, or the catalytically inactive D498N mutant, Addgene plasmid #259311) were generated using lentiviral transduction as previously described(*59*). Either parental, or ATP13A2 and ATP13A3 knockout cell lines were used. Following transduction, cells were selected with puromycin (2 µg/ml; InvivoGen). Expression was confirmed by western blotting as described below and polyclonal cell lines with comparable expression levels were used for downstream experiments. A detailed protocol is available at protocols.io (DOI:https://dx.doi.org/10.17504/protocols.io.bw57pg9n).

### 14C-labeled polyamine uptake

Uptake of ¹⁴C-labeled polyamines (putrescine, PUT; spermidine, SPD; and spermine, SPM) was measured in SH-SY5Y cells as described previously (*21*), with minor adaptations. Cells were seeded in 12-well plates and grown to 70% confluency. Cells were then treated with either 5 µM 14C-labeled polyamine (ARC-[14C]-PUT, [14C]-SPD or [14C]-SPM) or 5 µM 14C-labeled polyamines supplemented with 100 µM unlabelled polyamines (PUT, SPD, or SPM) serving as a competition control to assess non-specific uptake. After 30 min incubation at 37°C, the medium was aspirated and cells were washed twice with ice-cold DPBS (without Ca2+/Mg2+, Sigma). Cells were subsequently lysed by incubation in 200 µl RIPA buffer (Thermo Scientific) for 10 min at room temperature followed by scraping. Lysates were transferred to scintillation vials containing 7 ml EcoLite™ Liquid Scintillation Cocktail (MP Biomedicals) followed by an additional wash with 200 µl ice-cold DPBS (without Ca2+/Mg2+, Sigma). Radioactivity (counts per minute, CPM) was quantified using a Tri-Carb™ 4910TR V Liquid Scintillation Counter (Revvity).

For uptake of ¹⁴C-labeled polyamines in the HEK293T cells, a similar protocol was followed as above. Cells were seeded in 12-well plates and grown to 70% confluency. For difluoromethylornithine (DFMO, Kind gift of Dr. P. M. Woster (Medical University of South Carolina, Charleston, SC)) condition, cells were treated with 1.5mM DFMO for 24 h prior to sample collection. Cells were treated with 0.1 µM of 14C-labeled polyamines in medium pretreated with 1mM aminoguanidine (AG). After 30 min incubation at 37°C, medium was aspirated and cells were washed once with ice-cold DPBS (without Ca2+/Mg2+). A detailed protocol is available at protocols.io (DOI: https://dx.doi.org/10.17504/protocols.io.rm7vz97y5gx1/v1).

### BODIPY-polyamine uptake assay

BODIPY (BDP)-polyamine probes (Merck) were prepared at 5 mM in 0.1 M MOPS (pH 7.0, adjusted with KOH) and stored at −20°C (*21*). For characterization and AMXT 1501 (Aminex Therapeutics) experiments, HEK293T cells were seeded in 12-well plates to reach 70% confluency on the day of the experiment. The cells were incubated with 0.1 µM BDP-polyamines for 30 min at 37°C. For antizyme experiments, HEK293T cells were seeded in antibiotic-free medium in 6-well plates to reach 60% confluency on day of transfection. Cells were transiently transfected with 0.8 µg hAZ1-TwinStrep®-HA (NM_004152.3) or hAZ2-TwinStrep®-HA (NM_002537.3) using FuGENE® HD (Promega) at a 1:3 DNA:FuGENE® HD ratio and 10 min incubation prior to addition to the cells. 24 h post-transfection, the cells were treated with BDP-SPD or BDP-SPM (see figure legends for additional details) for 30 min at 37 °C. Medium was pre-treated with 1 mM AG (Merck) for 30 min prior to AMXT 1501 and BDP-polyamine addition. For DFMO condition, cells were treated with 1.5 mM DFMO for 20 h (for antizyme experiments) or 24 h (for characterization and AMXT 1501 experiments) prior to cell collection. For AMXT 1501 treatments (with and without DFMO), cells were treated for 2 h with 1 µM AMXT 1501, prior to BDP-polyamine treatment. Cells were harvested in DPBS (without Ca²⁺/Mg²⁺, Sigma), centrifuged at 450 x g for 5 min and then resuspended in FACS buffer (0.1% BSA, 2 mM EDTA in DPBS). The median/mean fluorescent intensity (MFI) of 10,000 events was recorded using the BD FACSymphony™ A5 cell Analyzer (BD Biosciences, RRID:SCR_022538) for the uptake experiments following antizyme transfection in HEK293T cells or the Cytek Aurora (Cytek® Biosciences) for characterization of uptake in HEK293T cells.

In SH-SY5Y cells, a similar protocol was followed as above for AMXT 1501/AMXT 1505 and BDP-polyamine treatments. The median/mean fluorescent intensity (MFI) of 10,000 events was recorded using the BD FACSymphony™ A5 cell Analyzer (BD Biosciences).

In M229/M229R cells, a similar protocol was followed as above with the minor modifications. Cells were treated with 1 µM BDP-polyamines. The median/mean fluorescent intensity (MFI) of 10,000 events was recorded using the BD FACS Canto™ (BD Biosciences). A detailed protocol is available at protocols.io (DOI: https://dx.doi.org/10.17504/protocols.io.eq2lymp3rlx9/v1).

### Polyamine toxicity rescue assay

#### 4-methylumbelliferyl heptanoate (MUH) assay

MUH assay was performed as previously described (*60*) (*16*), with minor modifications. SH-SY5Y ATP13A3 WT O.E. were seeded in 96-well plates to get 80% confluency cells on the day of the experiment. Cells were pretreated for 2 h with different concentrations of AMXT 1501 (medium pretreated with 1mM AG for 30 min). Subsequently, toxic concentrations of polyamines (PUT, SPD, and SPM) were added to the cells in the presence of AMXT 1501 and incubated overnight. MUH luminescence was measured using a BioTek Synergy H1 Multimode Reader (Agilent, RRID:SCR_019748) Data were controlled by subtracting the background fluorescence of positive control wells (toxic doses of polyamines; 0% viability) and expressed as a percentage relative to the negative control wells (untreated; 100% viability). A detailed protocol is available at protocols.io (doi.org/10.17504/protocols.io.bazjif4n).

#### CellTiter-Glo® luminescent cell viability assay

HEK293T ATP13A3 KO cells with stable overexpression of ATP13A3 WT were seeded in 384-well plates to get 80% confluency cells on the day of the experiment. 2 µg/ml Poly-D-Lysine (Gibco) was added to promote adhesion of the HEK cells. The cells were treated for 2 h with AMXT (medium pre-treated with 0.5 mM AG) followed by 0.1 mM SPD treatment for 24 h. Cell viability was measured using the CellTiter-Glo® luminescent cell viability assay (Promega) according to the manufacturer’s instructions. Luminescence was read out with a BioTek Synergy H1 Multimode Reader (Agilent, RRID:SCR_019748). Data were controlled by subtracting the background luminescence of wells containing 0% viable cells, expressed as a percentage relative to the untreated 100% viable controls.

#### Co-Immunoprecipitation (co-IP)

HEK293T ATP13A3 KO cells stably expressing ATP13A3 WT or ATP13A2 WT were seeded in 145 cm² dishes, in an antibiotic-free medium, to reach 60% confluency on the day of transfection. Cells were transiently transfected with 8 µg hAZ1-TwinStrep®-HA or hAZ2-Twin-Strep®-HA using FuGENE® HD (Promega) at a 1:3 DNA:FuGENE ® HD ratio and 10 min incubation prior to addition to the cells. To confirm the specific interaction between ATP13A3 and AZ1/AZ2, co-IP was also performed using non-transfected cells. Cells were harvested 24 h post-transfection and co-IP was performed using the ChromoTek® HA-Trap agarose kit (ProteinTech), according to the manufacturer’s instructions. Input and IP fraction were analyzed by SDS-PAGE and western blotting to detect ATP13A3, ATP13A2 and AZ1-HA or AZ2-HA as described below. A detailed protocol is available at protocols.io (DOI: https://dx.doi.org/10.17504/protocols.io.8epv5k3r5v1b/v1).

#### Protein expression in HEK293T cells

hATP13A2 (isoform 2, NM_001141973.3) and hATP13A3 (isoform 1, NM_024524.4) were cloned into the pcDNA3.4 vector with a C-terminal TwinStrep® and FLAG-tag. HEK293T cells (at 80% confluency and in a serum-, antibiotic-free medium) were transiently transfected with either ATP13A2 or ATP13A3 plasmids using polyethylenimine MAX (PEI MAX®; Polysciences) at a 1:4 DNA:PEI ratio. After 4 h at 37 °C, an equal volume of medium (containing serum and antibiotics) was added. Cells were harvested 72 h post-transfection by scraping in DPBS (without Ca2+/Mg2+, Sigma) containing 0.2% EDTA (VWR), centrifuged at 1,500 x g for 10 min, washed with ice-cold DPBS (without Ca2+/Mg2+, Sigma) and flash-frozen in liquid nitrogen before storing at −80°C until use. A detailed protocol is available at protocols.io (DOI: https://dx.doi.org/10.17504/protocols.io.36wgqpm5xvk5/v1).

#### Affinity purification from HEK293T cells

All steps were performed at 4°C unless stated otherwise. Frozen cell pellets were thawed, resuspended at 0.2 g wet weight per mL in hypotonic lysis buffer (10 mM Tris-HCl pH 7.5, 0.5 mM MgCl₂, 5 mM TCEP and SIGMAFAST™ protease inhibitor cocktail (Merck)) for 15 min and lysed with 60 strokes by Dounce homogenizer (DWK Life Sciences). Equal volume of the isotonic buffer (10 mM Tris-HCl pH 7.5, 0.5 M sucrose, 40 µM CaCl₂, 5 mM TCEP and SIGMAFAST™ protease inhibitor cocktail (Merck)) was added followed by an additional 40 strokes. Afterwards, the lysate was clarified at 2,000 x g for 20 min to remove nuclei and debris. Membrane solubilization was performed by adding n-dodecyl-β-D-maltopyranoside/Cholesteryl hemisuccinate (DDM/CHS; 10:1) at a 3:1 detergent-to-protein ratio for 2 h on a head-over-head rotator. Non-solubilized material was removed by centrifugation at 16,000 x g for 1 h. The supernatant was incubated with StrepTactin® XT resins (IBA Lifesciences) for 1 h and purified in batch mode in a gravity column (Econo-Pac®, BioRad). The resins were washed twice with 10 column volumes of each wash buffer (wash buffer 1: 20 mM Tris-HCl pH 7.5, 300 mM NaCl, 0.03% DDM, 0.003% CHS, 5 mM TCEP and wash buffer 2: wash buffer 1 with 150 mM NaCl). Protein was eluted using biotin-containing elution buffer (100 mM Tris-HCl pH 8.0, 150 mM NaCl, 1 mM EDTA, 50 mM biotin, 0.03% DDM, 0.003% CHS, 5 mM TCEP). Purification fractions were analyzed by SDS-PAGE with Coomassie staining (InstantBlue® Coomassie Protein stain, Abcam) or SYPRO Ruby staining (ThermoFisher) using BSA standard curve for concentration determination or western blotting as described below. A detailed protocol is available at protocols.io (DOI: https://dx.doi.org/10.17504/protocols.io.5qpvodq29g4o/v1).

#### Protein expression and purification from yeast

*S. cerevisiae* BY4743 (MATa/MATα; his3Δ1/his3Δ1, leu2Δ0/leu2Δ0, LYS2/lys2Δ0, met15Δ0/MET15, ura3Δ0/ura3Δ0, YPL217c/YPL217c::kanMX4; gift from J. Winderickx, KU Leuven) was transformed using the pYSG-IBA162 vector encoding a yeast codon-optimized hATP13A3 (isoform 1, NM_024524.4) with a C-terminal TwinStrep tag, as described for hATP13A2 (*20, 61*). Transformation was done using the lithium acetate/single-stranded carrier DNA/polyethylene glycol method and selected on MM-Ura agar (0.54% yeast nitrogen base without amino acids, 0.12% yeast drop-out mix without uracil, 2% glucose, 2% agar) at 30°C for 48 h. A single colony was expanded through sequential pre-cultures in MM-Ura liquid medium (0.67% yeast nitrogen base without amino acids, 0.19% yeast drop-out mix without uracil, 2% glucose) at 28°C, 175 rpm, and used to inoculate 4.5 L MM-Ura (starting OD₆₀₀ = 0.05) for overnight growth. Cells were harvested (1,000 x g, 10 min, 4°C), resuspended in 4.5 L MM-Leu medium (0.67% yeast nitrogen base without amino acids, 0.19% yeast drop-out mix without leucine, 2% glucose), and expression was induced with 0.5 mM CuSO₄ for 24 h. Cells were collected (1,000 x g, 10 min, 4°C) and lysed with glass beads (Sigma) in a BeadBeater (BioSpec Products) in lysis buffer: 50 mM Tris-HCl pH 7.5, 1 mM EDTA, 0.6 M sorbitol, 1 mM PMSF, SIGMAFAST™ protease inhibitor cocktail (Merck). The lysate was clarified at 2,000 × g for 20 min at 4°C (S1). S1 was centrifuged at 20,000 × g for 20 min at 4°C to obtain the heavy membrane fraction (P2). The resulting supernatant (S2) was ultracentrifuged at 200,000 × g for 1 h at 4°C to collect the light membrane fraction (P3), resuspended in 20 mM HEPES-Tris pH 7.4, 0.3 M sucrose, 0.1 mM CaCl₂. Protein concentration was determined by Bradford assay (Sigma). P3 membranes were diluted to 5 mg/mL in solubilization buffer (50 mM Tris-HCl pH 7.5, 150 mM NaCl, 20% glycerol, 5 mM dithiothreitol (DTT), 0.5% (wt/vol) DDM, SIGMAFAST™ protease inhibitor cocktail (Merck)) and incubated with gentle stirring on ice for 60 min. Insoluble material was removed by ultracentrifugation at 100,000 × g for 1 h at 4°C. The clarified supernatant was incubated with StrepTactin® XT 4Flow® resin (IBA Lifesciences) for 3 h at 4°C, washed four times with 10 column volumes of wash buffer (solubilization buffer with 0.03% DDM), and eluted with wash buffer supplemented with 50 mM biotin. A detailed protocol is available at protocols.io (doi.org/10.17504/protocols.io.ewov1d9jpvr2/v1).

#### Purification of N-terminal ATP13A3 peptide

Recombinant N-terminal ATP13A3 fragment (ATP13A3-NT), fused to GST, was expressed in *E. coli* BL21-AI cells using a pDEST15 vector. Bacterial pellets were lysed, inclusion bodies removed, and GST affinity purification performed according to the manufacturer’s instructions (Life Technologies). The purified protein was eluted in 50 mM Tris-HCl pH 8.3, 150 mM NaCl, and 50 mM reduced glutathione.

#### Protein expression and purification of human antizyme 2

Antizyme 2 (NM_002537.3) was expressed in the *E. coli* strain BL21 (DE3) using the pACYCDuet-1 vector encoding a N-terminal Histidine tagged construct. From a single colony, a seed culture of 20 ml was grown ON at 37 °C (200 rpm) in LB medium. This was diluted to an OD600 of around 0.1 into a 600 ml main culture and grown at 37 °C. Protein expression was induced at OD600 of 0.7-0.8 with 0.1 mM IPTG and incubated for 22-24 hours at 18 °C (200 rpm). The cells were collected by centrifugation at 16,000 xg for 10 min. For purification, the cell pellets were incubated with the lysis buffer: 40 mM Tris-HCl, pH 7.5, 300 mM NaCl, 5 mM β-mercaptoethanol, 0.1 mg/ml chicken egg white lysozyme (Merck), 4 µg/ml DNase I (Merck) and SIGMAFAST™ protease inhibitor cocktail (Merck) with 7 ml of buffer per 1 g of dry pellet. The incubation was performed with gentle shaking, first for 20 min at RT followed by 30 min at 4 °C. Cells were lysed using a probe sonicator with microtip (Branson) for 4 × 20 s at 60% amplitude, with 1 min intervals between pulses. Following centrifugation at 16,000 x g, the supernatant was added to TALON® Superflow™ metal affinity resin (Cytiva) (1 ml resin per 30 ml of supernatant) and incubated for 2 hrs at 4 °C in a head-over-head rotator. The resin washed with 3x column volume per wash using the wash buffer (20 mM Tris-HCl, pH 7.5, 300 mM NaCl) supplemented with sequentially increasing concentrations of imidazole (10 mM, 20 mM, and 30 mM imidazole for wash buffers 1, 2 and 3 respectively). Antizyme was eluted using wash buffer supplemented with 200 mM imidazole. Buffer exchange and concentration of the eluted protein into an imidazole-free buffer was performed using the Amicon® Ultra centrifugal filter 10 KDa MWCO (Millipore) according to the manufacturer’s instructions.

#### Lipid-protein overlay

Membrane strips spotted with lipids (Echelon™ biosciences) were blocked for 1 h at room temperature in buffer A (10 mM Tris-HCl pH 7.0, 150 mM NaCl, 0.1% Tween-20, 0.2% ovalbumin). Purified ATP13A3-NT fragment (5.3 nM final concentration) was then added and incubated for 1 h. After three 5-min washes with wash buffer (10 mM Tris-HCl pH 7.0, 150 mM NaCl, 0.1% Tween-20), membranes were probed for 1 h with anti-GST antibody (1:2500; 27-4577-01, RRID:AB_771432, Cytiva) diluted in buffer A. Strips were subsequently washed and incubated for 1 h with HRP-conjugated anti-goat antibody (1:2000, diluted in buffer A, sc-2020, RRID:AB_631728, Santa Cruz). Following three washes with wash buffer and an additional three washes with wash buffer lacking Tween-20, bound proteins were visualized using the ECL substrate (GE Healthcare) and imaged with a ChemiDoc™ imaging system (Bio-Rad).

#### Cell lysate preparation

Cell pellets were incubated for 30 min on ice in RIPA buffer supplemented with protease inhibitor cocktail SigmaFAST (S8830; Sigma). Lysates were centrifuged at 20,000 g for 35 min, and protein concentrations of the supernatants were determined using the BCA assay (Pierce, Thermo Fisher).

#### Western blotting

Samples (cell lysates or purification fractions) were mixed with LDS sample buffer (NP0008, ThermoFisher) containing 5 mM DTT. Proteins were separated on NuPAGE 4-12% Bis-Tris gels (Invitrogen) alongside the SeeBlue™ Plus2 molecular weight marker (LC5925, Thermo Scientific), using MOPS running buffer (NP0001, Thermo scientific) or MES running buffer (Thermo Scientific) for 1 h. Proteins were transferred onto PVDF membranes (1704157, Bio-Rad; IPFL00010, Millipore) by either semi-dry transfer using the Trans-Blot Turbo system (Bio-Rad; 30 min, 1.0 A, 25 V) or classical wet transfer (for yeast purification samples and visualisation of ATP13A3), according to the manufacturer’s protocols. Membranes were blocked in 5% non-fat milk or 3% BSA in tris-buffered saline (50 mM Tris-HCl, 150 mM NaCl) containing 0,1% tween 20 (TBS-T), then incubated overnight at 4°C with primary antibodies. Antibodies used included anti-ATP13A2 (1:2000, A28696, RRID:AB_3741517, ABclonal), anti-ATP13A3 (1:1000, HPA029471, RRID:AB_10600784, Sigma), anti-FLAG (1:1000, MA1- 91878, RRID:AB_1957945, Thermo Fisher), anti-streptavidin (1:1000, ab76949, RRID:AB_1524455, Abcam), anti-AZ2 (1:1000, PA5-103991, RRID:AB_2853322), anti-HA (1:1000, ab18181, RRID:AB_444303), anti-AZ1 (1:1000, PA5-77217, RRID:AB_2720944) and anti-α-tubulin (1:1000, T5168, RRID:AB_477579) diluted in TBS-T containing 1 or 3% BSA. After two washes in TBS-T, membranes were incubated for 1 h with HRP-conjugated secondary antibodies diluted in TBS-T containing 3% BSA: anti-mouse IgG (1:1000, 7076, RRID:AB_330924, Cell Signaling) and anti-rabbit IgG (1:1000, 7074, RRID:AB_2099233, Cell Signaling). Following final washes in TBS-T, bands were visualized using SuperSignal™West Pico PLUS chemiluminescent substrate (34580, Thermo Scientific) and imaged with a ChemiDoc™MPimaging system (Bio-Rad).

#### ATPase activity assay

ATPase activity was measured using the ADP-Glo™ Max luminescence assay (Promega) as previously described (*20*), with minor modifications. Reactions (25 µL) contained 80 nM purified hATP13A3, 11 mM MgCl₂, 100 mM KCl, 50 mM MOPS-KOH (pH 7.0), 1 mM DTT, 0.002% (wt/vol) DDM, 0.0002% (wt/vol) CHS, and the indicated polyamine concentrations. Reactions were initiated by addition of 2 mM ATP at 37°C for 30 min and terminated by addition of 25 µl ADP-Glo reagent, followed by 40 min incubation at room temperature. Subsequently, 50 µl of ADP-Glo Max Detection Reagent was added and incubated for 1 h at room temperature in the dark. To assess antizyme inhibition, the reaction volume was halved to 12.5 µL and hAZ2 was added at different molar ratio with hATP13A3 (80 nM each). Luminescence was measured on a Synergy H1 microplate reader (BioTek). Kinetic parameters were calculated by curve fitting in GraphPad Prism. A detailed protocol is available in protocols.io. DOI: https://dx.doi.org/10.17504/protocols.io.8epv5k3zdv1b/v1.

#### Auto-phosphorylation assay

The ability of ATP13A3 to form a phosphorylated intermediate at the conserved D498 residue was examined using radiolabelled ATP, following established procedures (*20*) (*37*) with minor modifications. P2 yeast membranes (40 μg protein) or purified hATP13A3 (1 μg) were incubated with [γ-32P] ATP (2 μCi, 5.125 µM, Revvity) for 1 min at 4°C in the absence or presence of indicated polyamines (1 mM). To evaluate nucleotide sensitivity, either non-radioactive ADP or ATP (5 mM each) was added 30 s after [γ-32P] ATP, and reactions were terminated at the indicated time points. For combined treatments, non-radioactive ATP (5 mM) and polyamines (1 mM) were added simultaneously 30 s post [γ-32P] ATP addition. For AZ2 inhibition experiments, purified hATP13A3 was pre-incubated with purified hAZ2, at varying molar ratios, for 30 min at 4°C prior to initiating the assay. Reactions were stopped by addition of 400 μL ice-cold stop solution (20% trichloroacetic acid, 10 mM phosphoric acid) and incubated on ice for 30 min. Precipitated material was pelleted by centrifugation (20,000 x g, 30 min, 4 °C), washed twice with ice-cold stop solution and dissolved in sample buffer 1 for ATP13A3 characterization experiments: (5mM NaH2PO4, pH 6, 0.005% SDS, 0.1 g/ml LDS, 10% glycerol, 0.5 mg/ml bromophenol blue, 20 µl/ml β-mercaptoethanol) or sample buffer 2 for ATP13A3-AZ2 experiments: 50 mM MOPS, pH 6.5, 10% glycerol, 10 mM DTT, 5% SDS, 0.5 mg/ml bromophenol blue. Where indicated, the pellets received an additional wash in 0.3 M hydroxylamine (HA) prior to resuspension. Samples were resolved by SDS-PAGE under acidic conditions, and incorporation of 32P was detected by autoradiography using a Typhoon™ FLA 7000 (GE Healthcare). A detailed protocol is available at protocols.io. (DOI: https://dx.doi.org/10.17504/protocols.io.ewov1e8xogr2/v1)

#### RT-qPCR

RNA was isolated from 1.0 × 10⁶ cells using the NucleoSpin® RNA plus kit (Macherey-Nagel) following manufacturer’s instructions. RNA concentration and purity were determined using a Nanodrop 1000 Spectrophotometer (Thermo Scientific™, RRID:SCR_016517). cDNA was synthesized from 2 µg RNA using the High-Capacity cDNA Reverse Transcription Kit (Applied Biosystems™) according to the manufacturer’s instructions. For quantitative PCR, cDNA samples were diluted 1:25, and 2.5 µL of diluted cDNA was loaded per well in duplicate. The following reaction mixture was added to each well (final volume 10 µL): 5 µL PowerUp SYBR™ Green Master Mix (2×; Applied Biosystems™), 0.5 µL forward primer (10 µM), 0.5 µL reverse primer (10 µM), and 1.5 µL Milli-Q water. Negative control reactions were performed by replacing cDNA with an equal volume of Milli-Q water. PCR amplification was carried out in fast cycling mode with the following conditions: UDG activation at 50°C for 2 min, dual-lock DNA polymerase activation at 95°C for 2 min, followed by 40 cycles of denaturation at 95°C for 1 s (QuantStudio™ 5, Applied Biosystems™, RRID:SCR_020240) or 3 s (StepOnePlus™, Applied Biosystems™, RRID:SCR_015805), and annealing/extension at 60°C for 30 s. Melt curves were generated using the step-and-hold method with a ramp rate of 0.4°/second to verify amplification specificity. PCR efficiencies were determined prior to relative expression analysis. Gene expression levels were normalized using GAPDH and/or ACTB as reference genes. The relative expression level was calculated using the comparative (2–ΔΔCT) method. The following primer sequences **(Table 2)** were utilized for RT-qPCR. A detailed protocol is available at protocols.io. DOI: https://dx.doi.org/10.17504/protocols.io.5qpvojyn7g4o/v1.

**Table 2:**
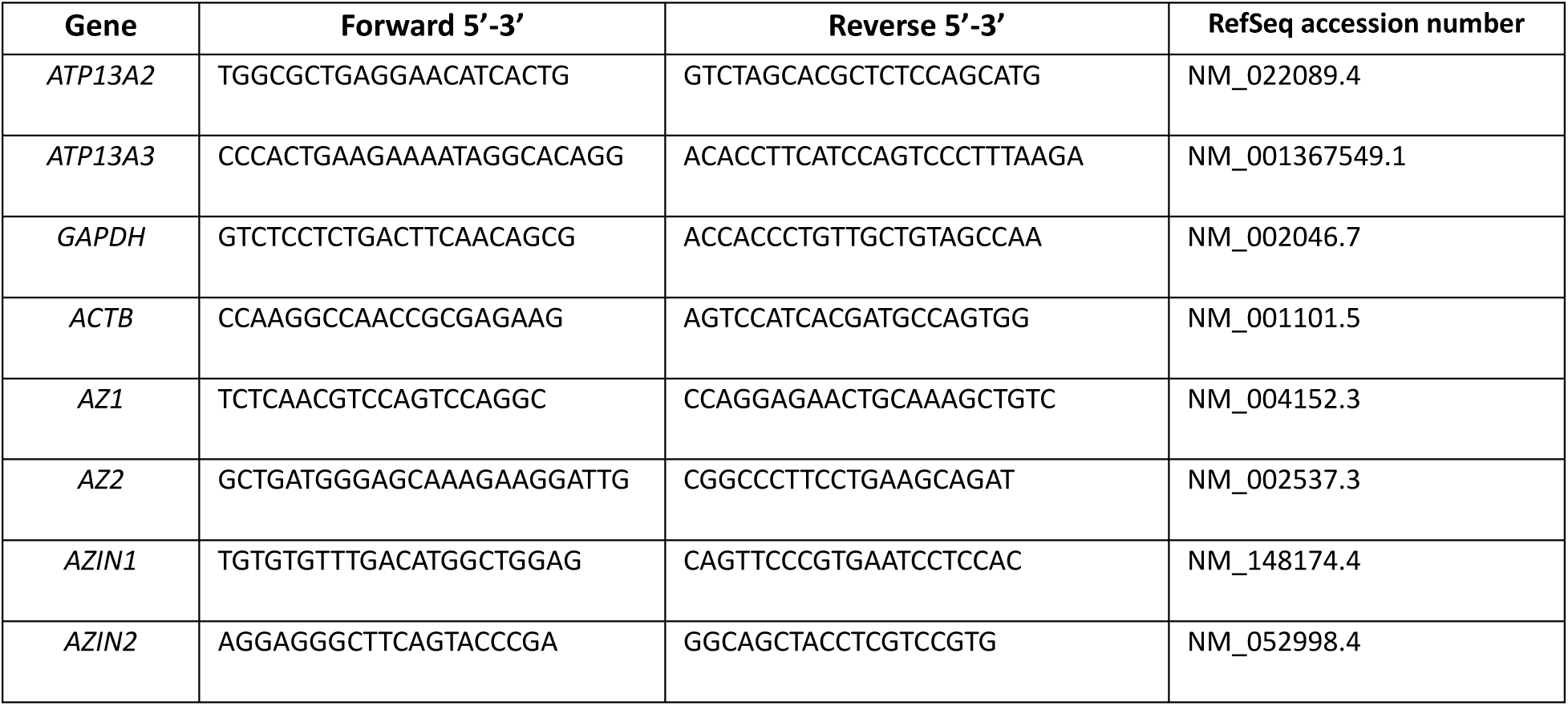
Primer used for RT-qPCRw.

#### AlphaFold prediction of protein-protein interactions

Three-dimensional structures of ATP13A3 (isoform 1), ATP13A2 (isoform 2), and AZ1-3 and their interactions were predicted using AlphaFold 3 (*36*). Input sequences were provided in single-letter amino acid format together with Mg²⁺ and ATP as cofactors. Model quality was assessed by predicted aligned error (PAE), predicted template modeling score (pTM), and interface pTM (ipTM). Only models with low PAE at the binding interface and ipTM ≥ 0.6 were used for downstream structural analysis in PyMOL (RRID:SCR_000305).

#### Multiple sequence alignment of antizyme proteins

Amino acid sequences of AZ1, AZ2, and AZ3 were aligned using MUSCLE (RRID:SCR_011812) and visualized using Jalview (*62*).

#### In silico analysis

Expression levels (log2-transformed) of ATP13A2, ATP13A3, OAZ1-2, ODC1 and AZIN1 were evaluated in primary and metastatic samples from The Cancer Genome Atlas - Skin Cutaneous Melanoma (TCGA-SKCM) cohort (*63*), accessed through the UCSC Xena platform (https://xena.ucsc.edu/, accessed on 24 June 2026) and the Gene Expression Omnibus (GEO) (*64*) GSE46517 dataset (*65*). For microarray data, one probe per gene was selected using the jetset R package (*66*). Groups were compared by Welch t-test with Benjamini-Hochberg correction. All data processing and statistical analysis were performed in R (version 4.4.3).

Tumor RNA-seq datasets were retrieved from The Cancer Genome Atlas (UCSC Toil RNA-seq Recompute Compendium (*67*)). 12 common TCGA cancer types were selected by tumor sample size for analysis to ensure adequate statistical power across all downstream analyses (differential expression and co-expression). All selected cancer types had ≥400 primary tumor samples. Normal tissue controls were assembled from GTEx normal tissue matched by primary anatomical site and TCGA matched adjacent normal tissue. The 12 cancer types were: breast invasive carcinoma (BRCA, n=1,099; matched controls, n=28), kidney renal clear cell carcinoma (KIRC, n=531; matched controls, n=100), brain lower grade glioma (LGG, n=523; matched controls, n=1152), head and neck squamous cell carcinoma (HNSC, n=520; matched controls, n=99), lung adenocarcinoma (LUAD, n=515; matched controls, n=247), thyroid carcinoma (THCA, n=512; matched controls, n=338), lung squamous cell carcinoma (LUSC, n=498; matched controls, n=338), prostate adenocarcinoma (PRAD, n=496; matched controls, n=152), skin cutaneous melanoma (SKCM, n=469; matched controls, n=813), ovarian serous cystadenocarcinoma (OV, n=427; matched controls, n=88), stomach adenocarcinoma (STAD, n=414; matched controls, n=210), and bladder urothelial carcinoma (BLCA, n=407; matched controls, n = 28). Co-expression between the six polyamine pathway genes (ATP13A2, ATP13A3, AZIN1, OAZ1, OAZ2, ODC1) was assessed using Spearman rank correlation within tumor samples of each of the 12 TCGA cancer types. All 15 unique gene pairs were tested per cancer type (180 tests total). Expression values were log2(FPKM+1) from the UCSC Xena Toil RNA-seq Recompute pipeline (*67*). Benjamini-Hochberg false discovery rate correction was applied across all 180 tests. Differential expression between tumor and matched normal tissue was assessed for the six polyamine pathway genes (ATP13A2, ATP13A3, AZIN1, OAZ1, OAZ2, ODC1) in the 12 TCGA cancer types. Expression values were log2(FPKM) from the UCSC Xena Toil RNA-seq Recompute pipeline. The log2 fold change (log2FC) was calculated as the difference in arithmetic means between tumor and normal samples in log2 space [log2FC = mean(log2 FPKM)_tumor − mean(log2 FPKM)_normal]. The 95% confidence interval for the log2FC was derived from the standard error of the difference between two independent group means [SE(log2FC) = √(SE²_tumor + SE²_normal)]. Statistical significance was assessed using the two-sided Wilcoxon rank-sum test (Mann-Whitney U test; SciPy v1.15.0). Benjamini-Hochberg false discovery rate (BH-FDR) correction was applied across all tests within each analysis (72 tests for the 6-gene analysis).

Analysis of gene expression during drug treatment was assessed in a melanoma model of a single BRAF V600E patient-derived xenograft model (MEL006) treated with dabrafenib and trametinib combination therapy (taken from GEO dataset GSE116237(*68*)). Single-cell RNA-seq count data were normalized to 10,000 counts per cell and log1p-transformed using the Scanpy framework (*69*). For each of the six polyamine pathway genes (*ATP13A2*, *ATP13A3*, *AZIN1*, *OAZ1*, *OAZ2*, *ODC1*), per-cell normalized expression values were compared across four progression groups defined by the experimental design: T0 (pre-treatment, n = 172 cells), T4 (4 days on treatment, n = 155), T28 (minimal residual disease at 28 days, n = 99), AR (acquired resistance, n = 100). Differential expression between groups was assessed using the two-sided Mann-Whitney U test (Wilcoxon rank-sum test), applied per gene to the per-cell log-normalized expression values. An omnibus Kruskal-Wallis test was additionally performed for each gene across all four groups. Effect sizes were quantified as the difference in mean log-normalized expression between groups (log2FC) and as the area under the receiver operating characteristic curve (AUC), derived from the Mann-Whitney U statistic as AUC = U / (n₁ × n₂), where n₁ and n₂ are the group sizes. Multiple-testing correction was applied using the Benjamini-Hochberg procedure to control the false discovery rate (FDR) at 5%, with correction applied across the seven genes within each pairwise comparison. Genes with adjusted p-values (padj) below 0.05 were considered statistically significant. Individual cells were treated as independent replicates for statistical testing, consistent with standard single-cell analysis conventions. Data processing and analyses of the TCGA-derived datasets and the GEO dataset GSE116237 were performed in Python using SciPy(*70*) and statsmodels (*71*).

#### Statistical analysis

Statistical analyses were performed using GraphPad Prism version 10 (RRID:SCR_002798). Statistical analysis of *in silico* analysis of gene expression is described under *in silico* analysis. The specific statistical tests applied are indicated in the respective figure\ legends. Data are presented as mean ± standard error of the mean (SEM), and individual data points are shown in bar graphs, where applicable. All experiments were performed with at least three independent biological replicates.

