## Supplementary Figures for "A druggable ATP13A3–antizyme switch controls adaptive polyamine uptake in cancer"

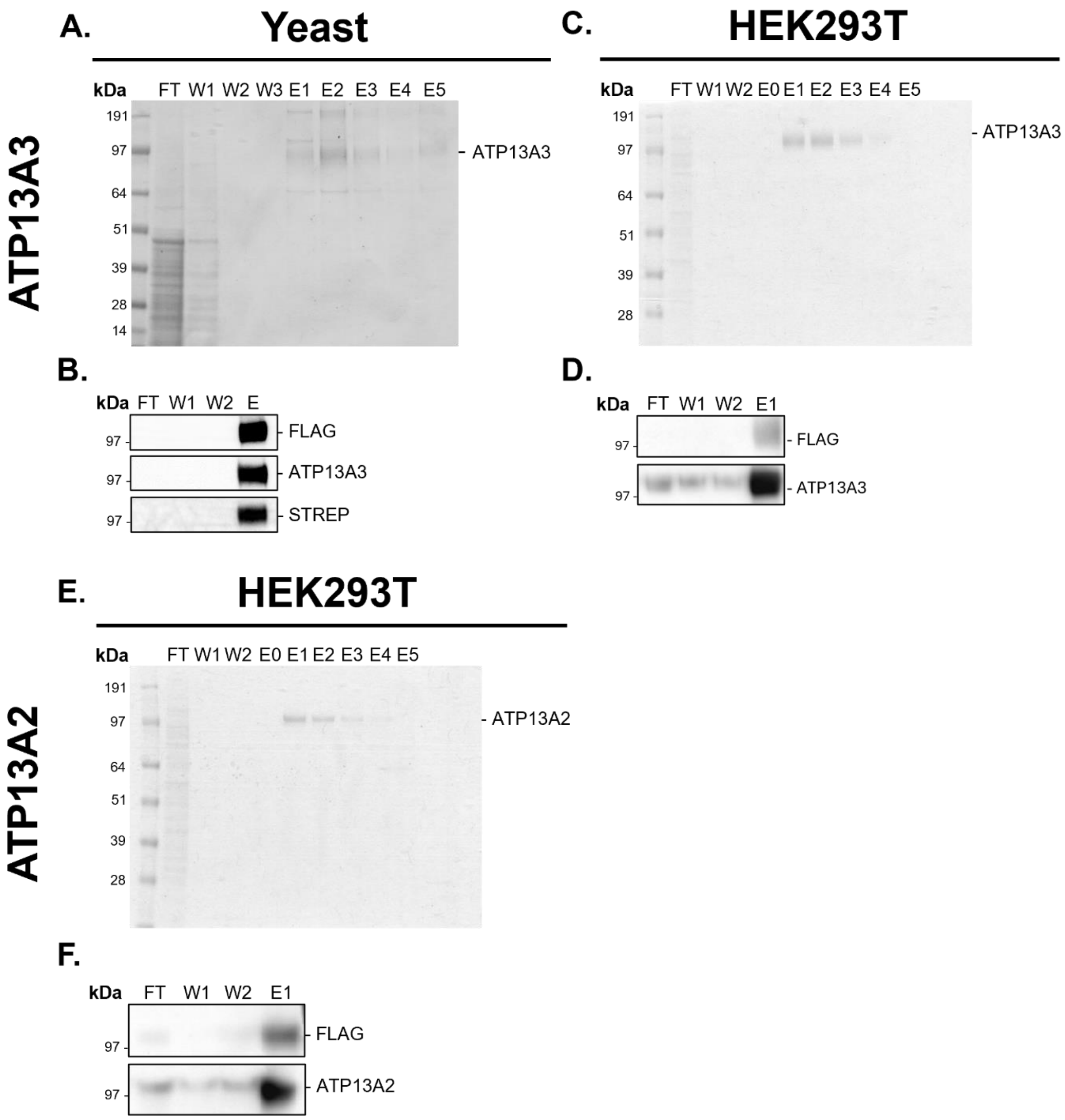


**Figure S1: Purification of hATP13A3 and hATP13A2 from yeast and HEK293T cells.** Purification of FLAG/Strep-tagged hATP13A3 **(A-D)** and hATP13A2 **(E-F)** from *Saccharomyces cerevisiae* **(A-B)** and HEK293T cells **(C-F)**. **(A**, **C**, **E)** Coomassie staining and **(B**, **D**, **F)** Western blot analysis showing the purification process for WT hATP13A3 **(B, D)** and hATP13A2 **(F)** starting from detergent-solubilized yeast membranes **(A-B)** or HEK293T cell lysates **(C-F)**, followed by streptavidin affinity chromatography and subsequent elution with biotin. n $\geq$ 3 independent purifications, including flow-through (FT); wash (W1-W2(-W3)); elution ((E0)-E1-E5)

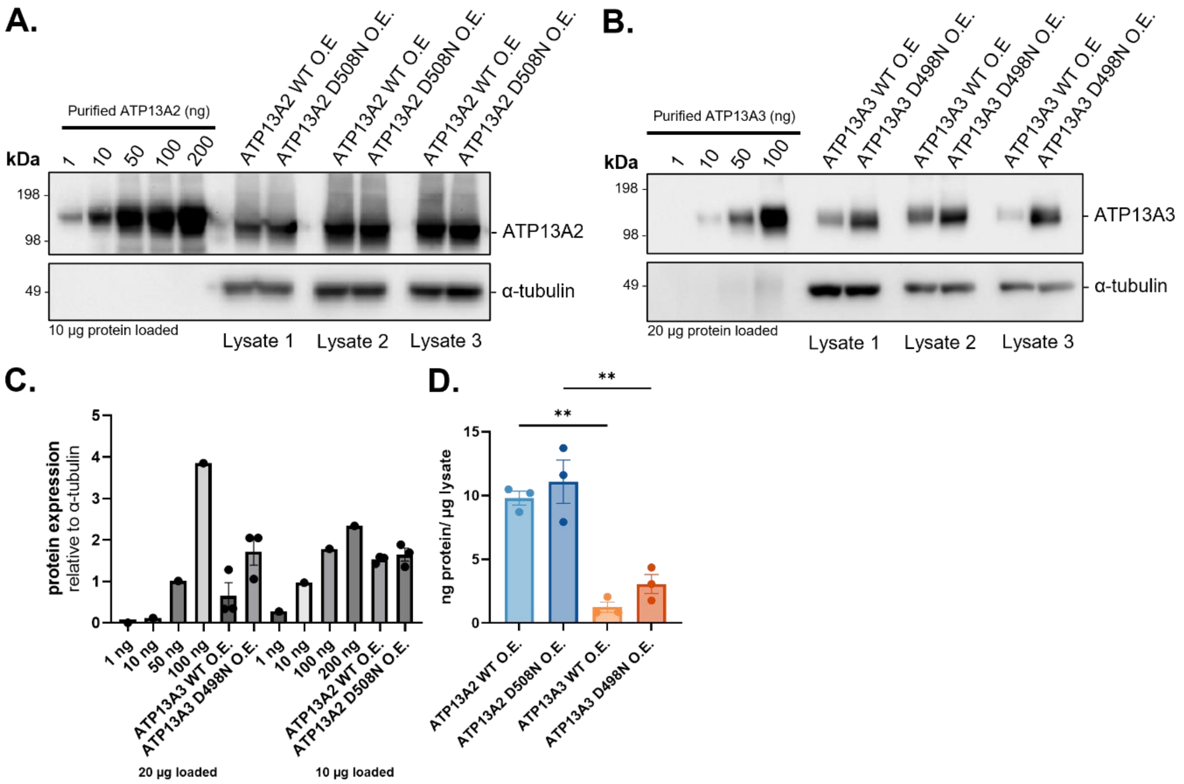


**Figure S2: Quantification of ATP13A2 and ATP13A3 protein expression levels in overexpression cell models. (A-B)** Immunoblots of purified human ATP13A2 (**A**; 1-200 ng) and human ATP13A3 (**B**; 1-100 ng) protein standards alongside lysates from SH-SY5Y cells stably expressing ATP13A2 WT or D508N, or ATP13A3 WT or D498N, respectively. These standards were used to generate standard curves for absolute quantification of ATP13A2 and ATP13A3 protein expression levels in cell lysates. α-Tubulin served as a loading control. Three independent lysates are shown for each cell line. **(C)** Quantification of ATP13A2 and ATP13A3 immunoblot signals shown in **(A)** and **(B)**. **(D)** Quantification of ATP13A2 and ATP13A3 protein concentration (ng protein per µg total lysate protein) in SH-SY5Y overexpression cell lines. Data are presented as mean ± SEM of *n* = 3 independent biological replicates, with individual data points shown. One-way ANOVA with Tukey's multiple-comparisons test.

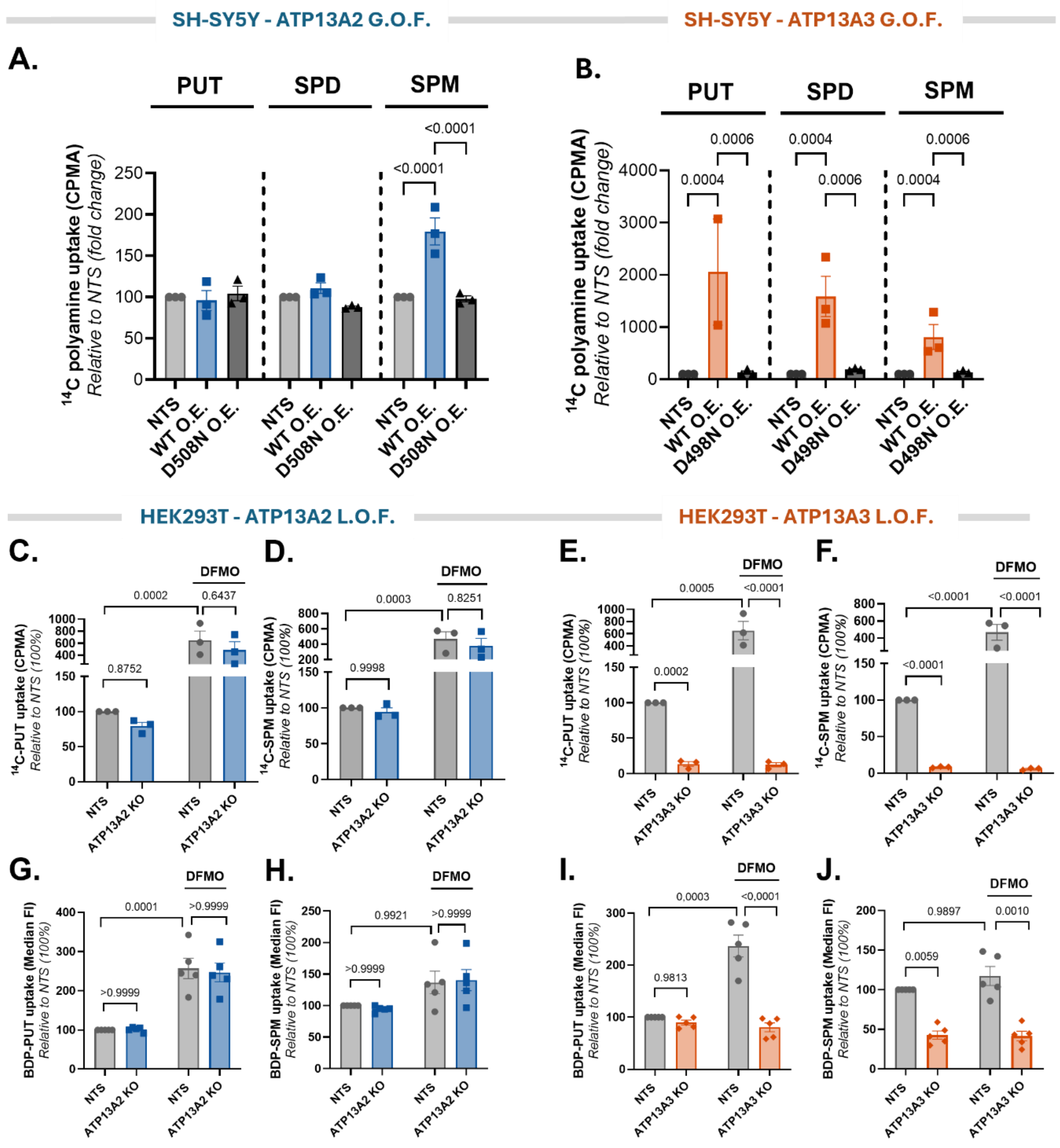


**Figure S3: Comparison of ATP13A2- and ATP13A3-mediated polyamine uptake in SH-SY5Y and HEK293T models.** Measurement of cellular polyamine uptake in ATP13A2 and ATP13A3 SH-SY5Y gain-of-function (G.O.F.) overexpression models and HEK293T loss-of-function (L.O.F.) knockout (KO) models. Uptake of $^{14}$C-radiolabeled putrescine (PUT), spermidine (SPD), and spermine (SPM) (5 µM, 30 min) in SH-SY5Y cells stably overexpressing ATP13A2 WT or D508N **(A)**, or ATP13A3 WT or D498N **(B)**. Uptake assays were performed in the presence of aminoguanidine to prevent extracellular polyamine oxidation. Uptake was normalized to non-transduced (NTS) cells for each polyamine. Data are presented as mean ± SEM of $n$ = 3 independent biological replicates, with individual data points shown. Statistical analysis was performed using two-way ANOVA followed by Tukey's multiple-comparisons test. Uptake of $^{14}$C-labelled PUT and SPM (1 µM, 30 min) **(C-F)** and BODIPY-labelled PUT and SPM (BDP-PUT and BDP-SPM; 0.1 µM, 30 min) **(G-J)** in HEK293T NTS, ATP13A2 KO **(C,D,G,H)** or ATP13A3 KO **(E,F,I,J)** cells in the presence or absence of 24h pre-treatment with 1.5 mM DFMO. Radiolabelled uptake was quantified by counts per minute (CPMA), whereas BODIPY-labelled polyamine uptake was quantified as median fluorescence intensity (MFI). Data are presented as uptake normalised to untreated NTS (set to 100%), with individual biological replicate values shown as dots and bars representing the mean ± SEM of $n$ = 3 or 5. Statistical analysis were performed on $\log_{10}$-transformed raw uptake values using two-way ANOVA followed by Šidák's post hoc multiple comparisons test. All experiments were performed with 1mM Aminoguanidine.

**A.**

| | Identity to ATP13A3 | Similarity to ATP13A3 |
|---|---|---|
| **ATP13A2** | 41.1 % | 58.7 % |
| **ATP13A4** | 44.9 % | 63.4 % |
| **ATP13A5** | 41.7 % | 61.6 % |

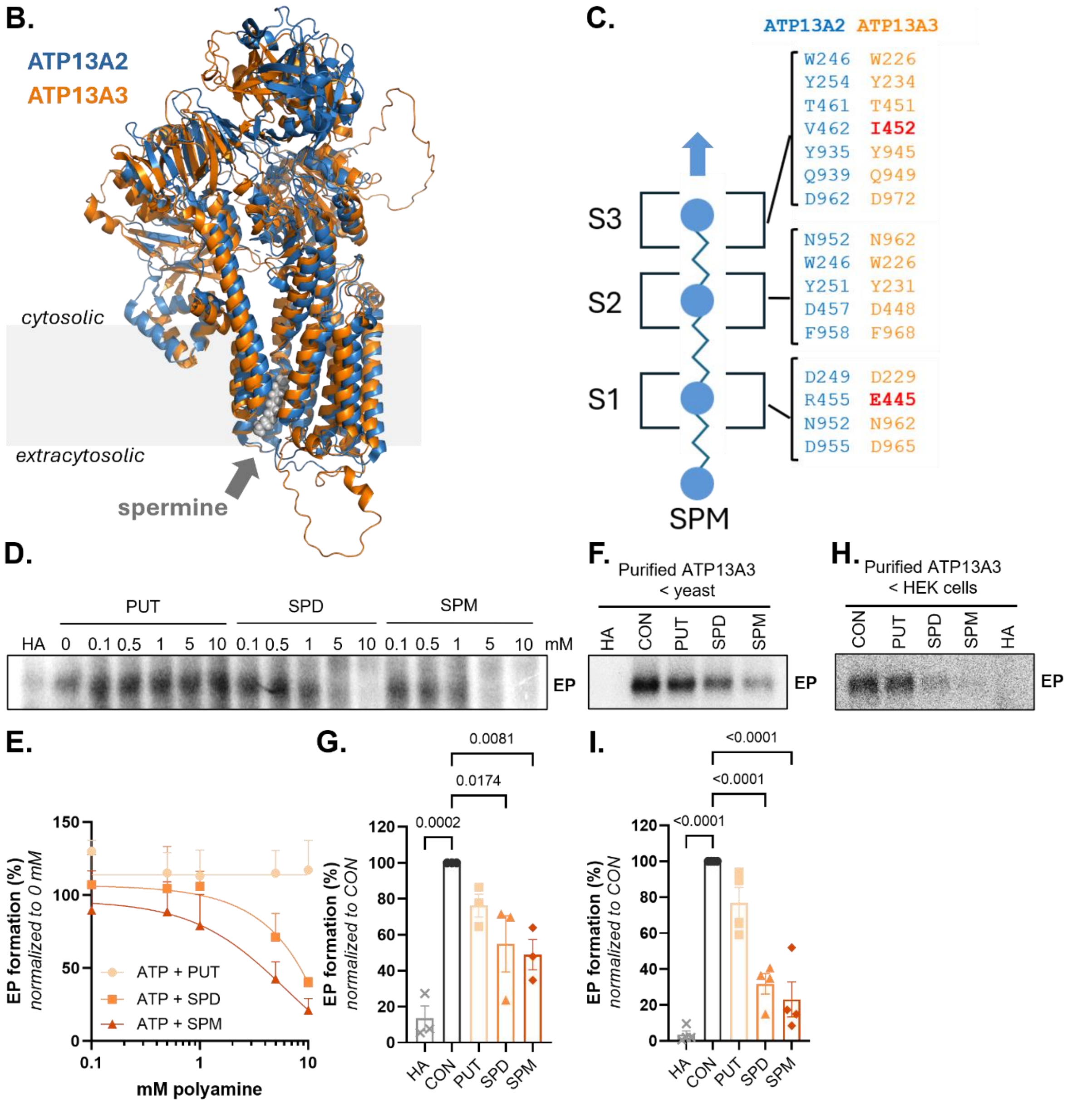


**Figure S4: Conservation of the polyamine-binding cavity between ATP13A2 and ATP13A3.** **(A)** Pairwise sequence identity and similarity of ATP13A2, ATP13A4, and ATP13A5 relative to ATP13A3. **(B)** Structural superposition cryo-EM structure of ATP13A2 in the spermine bound E2-Pi state (blue) PDB: 7n78 (*72*) and ATP13A3 alphafold prediction (orange) highlighting the conserved overall architecture of the transmembrane domain and the location of the polyamine-binding cavity (a bound spermine is indicated as grey spheres). **(C)** Comparison of residues lining the ATP13A2 and ATP13A3 polyamine-binding cavity. Residues forming the S2 coordination site are fully conserved between ATP13A2 and ATP13A3, whereas conservative amino acids substitutions are present at the S1 and S3 sites that may contribute to modest differences in polyamine affinity. **(D-E)** Representative autoradiogram **(D)** and quantification **(E)** of pulse ([γ-$^{32}$P]-ATP) chase (cold ATP) experiments in the presence of indicated concentrations of putrescine (PUT), spermidine (SPD), or spermine (SPM). Data are presented as mean ± SEM of $n$ = 2 independent biological replicates. **(F-I)** ATP13A3 phosphoenzyme levels in control (CON) conditions and in the presence of 1 mM putrescine (PUT), spermidine (SPD), spermine (SPM), or hydroxylamine (HA). **(F-G)** *S. cerevisiae* P2 membrane fractions with ATP13A3 expression. **(H-I)** ATP13A3 purified from HEK293T cells. **(F, H)** Representative autoradiograms. **(G, I)** Quantification of EP levels. Data are presented as mean ± SEM of $n$ = 3 **(G)** or $n$ = 4 (**I**) independent biological replicates, with individual data points shown. One-way ANOVA with Dunnett's multiple-comparisons test.

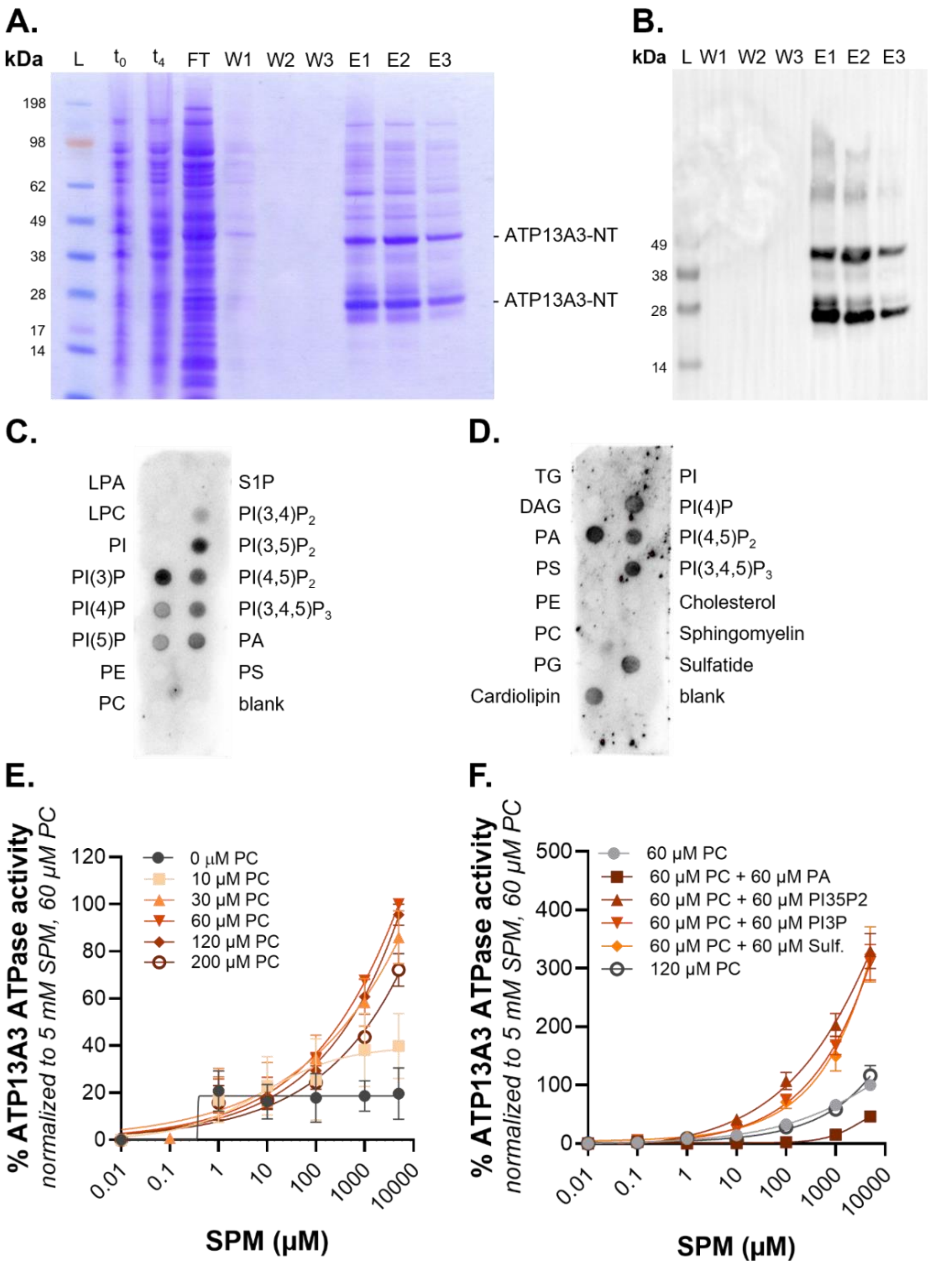


**Figure S5: Identification of ATP13A3 N-terminal lipid interactions and their impact on ATP13A3 ATPase activity. (A-B)** Purification of ATP13A3 N-terminal domain (ATP13A3-NT). **(A)** Coomassie-stained gel and **(B)** anti-His immunoblot showing samples collected during purification, including SeeBlue™ Plus2 Pre-stained Protein Standard (L); total lysates before ($t_0$) and after ($t_4$) induction; flow-through (FT); wash (W1–W3); and elution (E1–E3). **(C-D)** Lipid overlay assays of purified ATP13A3-NT using phosphoinositide containing **(C)** and membrane lipid-containing **(D)** strips. Bound ATP13A3-NT was detected by immunoblotting. Lipids tested included lysophosphatidic acid (LPA), lysophosphatidylcholine (LPC), phosphatidylinositol (PI), phosphatidylinositol phosphates (PIPs), phosphatidic acid (PA), phosphatidylserine (PS), phosphatidylethanolamine (PE), phosphatidylcholine (PC), phosphatidylglycerol (PG), diacylglycerol (DAG), triacylglycerol (TG), cardiolipin, cholesterol, sphingomyelin, and sulfatide. **(E-F)** Spermine (SPM)-stimulated ATPase activity of purified ATP13A3 measured in the presence of increasing concentrations of PC **(E)** or in the presence of 60 µM PC alone or supplemented with the indicated N-terminal interacting lipids **(F)**. ATPase activity was normalized to the activity measured in the presence of 5 mM SPM and 60 µM PC. Data are presented as mean ± SEM of $n$ = 3 **(E)** or 6 **(F)** independent biological replicates.

**A.**

| | ATP13A2 | | ATP13A3 | |
|---|---|---|---|---|
| | pTM | iPT M | pTM | iPT M |
| **AZ1** | 0.69 | 0.55 | 0.72 | 0.67 |
| **AZ2** | 0.72 | 0.58 | 0.73 | 0.77 |
| **AZ3** | 0.69 | 0.54 | 0.71 | 0.68 |

**B.**

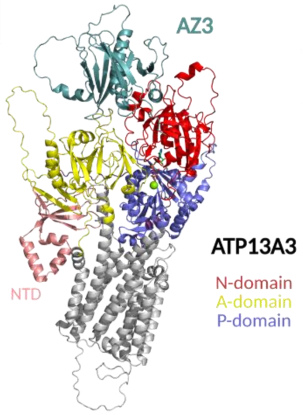


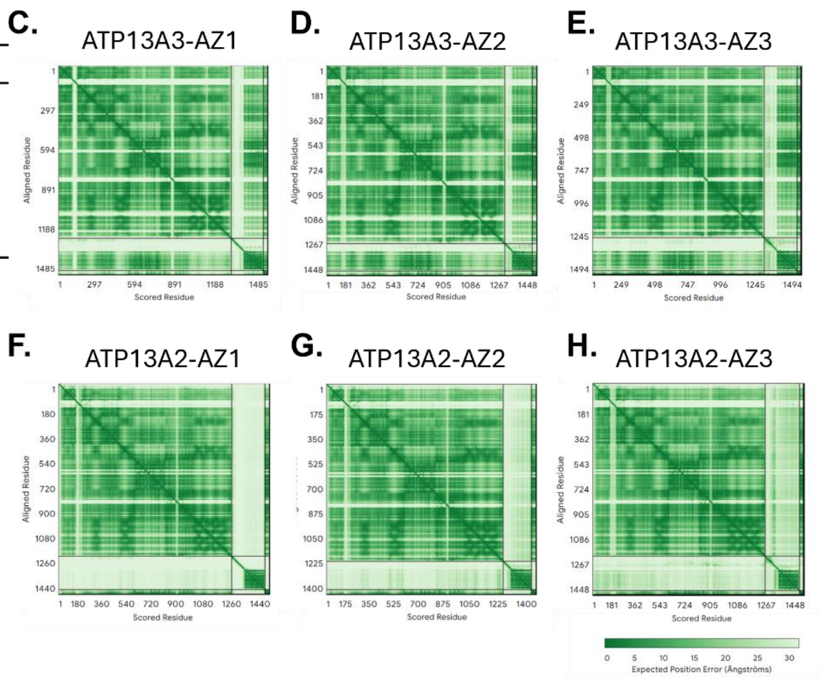


**I.**

| | Identity to AZ1 | Similarity to AZ1 |
|---|---|---|
| **AZ2** | 56.9 % | 70.7 % |
| **AZ3** | 36.3 % | 53.1 % |
| | **Identity to AZ2** | **Similarity to AZ2** |
| **AZ3** | 37.6 % | 56.1 % |

**Figure S6: Structural comparison of ATP13A2/ATP13A3 interactions with antizyme family members. (A)** AlphaFold heterodimer model confidence scores (pTM and ipTM) for predicted complexes between ATP13A2 or ATP13A3 and antizyme 1 (AZ1), antizyme 2 (AZ2), or antizyme 3 (AZ3). **(B)** Representative AlphaFold heterodimer model of the ATP13A3-AZ3 complex. ATP13A3 domains are colored as follows: N-domain (red), P-domain (purple), A-domain (yellow), transmembrane domain (grey), and N-terminal domains (NTD, pink). AZ3 is shown in cyan. **(C-E)** Predicted aligned error (PAE) plots for ATP13A3 in complex with AZ1 **(C)**, AZ2 **(D)**, or AZ3 **(E)**. **(F-H)** PAE plots for ATP13A2 in complex with AZ1 **(F)**, AZ2 **(G)**, or AZ3 **(H)**. Lower PAE values indicate higher confidence in the relative positioning of residues within the predicted complexes. **(I)** Pairwise sequence identity and similarity between AZ1, AZ2, and AZ3.

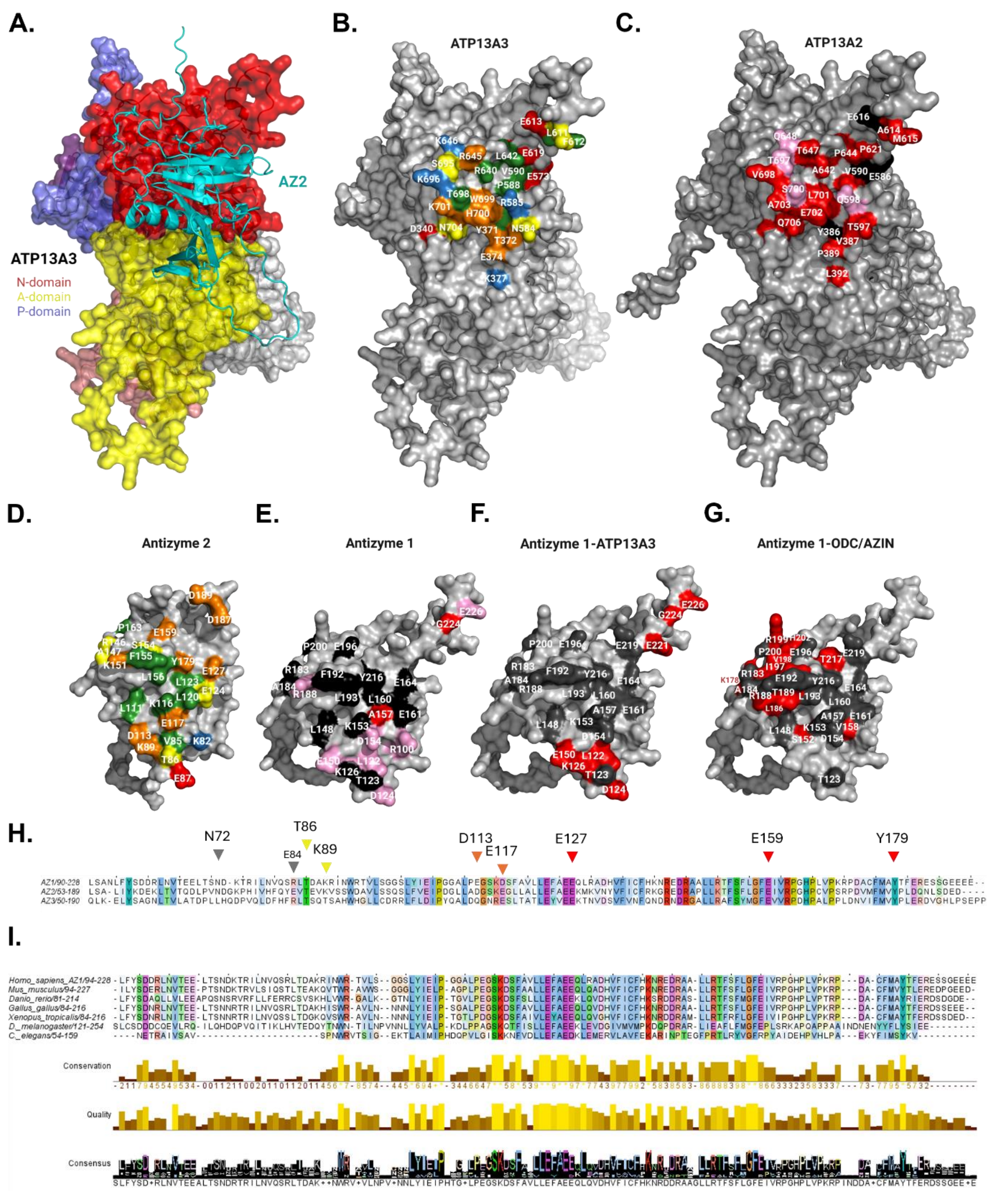


**Figure S7: Structural comparison of ATP13A2/ATP13A3 interactions with antizyme family members. (A)** Top view of the AlphaFold multimer model of the ATP13A3-AZ2 complex. ATP13A3 domains are colored as follows: N-domain (red), P-domain (purple), A-domain (yellow), transmembrane domain (grey), and N-terminal domains (NTD, pink). AZ2 is shown in cyan. **(B-C)** The AZ2-interacting surface of ATP13A3 differs significantly from corresponding residues in ATP13A2. **(B)** The predicted ATP13A3–AZ2 interaction surface is shown in open book view, with interface residues being colored according to their interaction types: green (van der Waals interaction), yellow (hydrogen bonding), red/blue (ionic interaction), and orange (involving two or more types of interactions or interaction partners). **(C)** Conservation of the ATP13A3 residues involved in the corresponding AZ2 interaction site on ATP13A2 are colored according to sequence conservation compared with ATP13A3: identical (black), similar (pink), and non-conserved (red).

**(D)** The ATP13A3-interacting surface of AZ2 is shown in open book view, with interface residues being colored according to their interaction types as done in panel (B) **(E)** Conservation of the AZ2 residues involved in the predicted ATP13A3 interaction for AZ1 re colored according to sequence conservation compared with AZ2 as done in panel (C) **(F-G)** Overview of the predicted ATP13A3 interacting residues of AZ1 and how they overlap with the interaction surface between AZ1 and ODC1/AZIN. Overlapping residues are coloured black while non-overlapping residues are coloured red. Residues involved in the AZ1-ODC1 and AZ1-AZIN interaction reported in (*7*) (PDB: 4ZGY and 4ZGZ) and their overlap with the AZ1-ATP13A3 interface. **(H)** Multiple sequence alignment of antizyme proteins. Residues are colored according to the ClustalX color scheme for visualization of conserved amino acids and physicochemical properties. Selected AZ2-residues for mutagenesis are indicated above the alignment. **(I)** Conservation of AZ1 across species. Multiple sequence alignment of AZ1 (OAZ1) protein orthologs from *Homo sapiens, Mus musculus, Gallus gallus, Xenopus laevis, Danio rerio, Drosophila melanogaster*, and *Caenorhabditis elegans*. The conservation and quality histograms indicate residue conservation and amino acid similarity across alignment positions, respectively. The consensus sequence is displayed below the alignment.

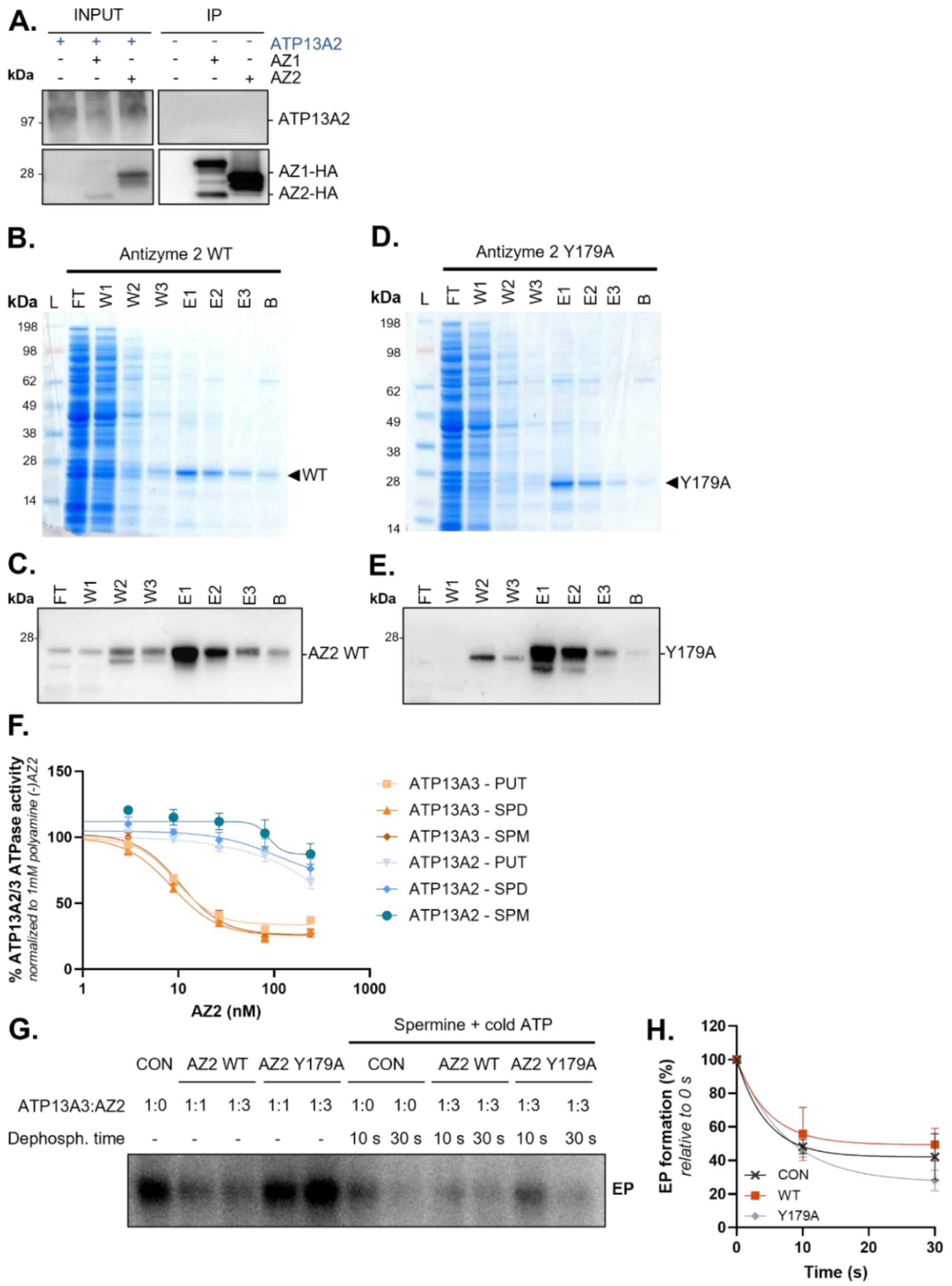


**Figure S8: Antizyme 2 WT and Y179A mutant purification and biochemical validation. (A)** Co-immunoprecipitation of ATP13A2 with AZ1-HA or AZ2-HA. Representative immunoblots of input and IP fractions. **(B-E)** Purification of antizyme 2

(AZ2) WT **(B, C)** and Y179A **(D, E)**. **(B, D)** Coomassie-stained gel and **(C, E)** Western blot analysis showing the purification process for AZ2 starting from lysed bacterial cells, followed by histidine-tag purification and subsequent elution with imidazole, including SeeBlue™ Plus2 Pre-stained Protein Standard (L); flow-through (FT); wash (W1-W3); elution (E1-E3); beads (B). N=3 independent purifications. **(F)** Putrescine (PUT), spermidine (SPD), and spermine (SPM)-stimulated ATPase activity of purified ATP13A3 and ATP13A2 measured in the presence of increasing concentrations purified AZ2. ATPase activity was normalized to the activity measured in the presence of 1 mM SPM. Data are presented as mean ± SEM of *n* = 3 independent biological replicates. **(G-H)** Effect of AZ2 on ATP13A3 phosphoenzyme (EP) formation. Representative autoradiogram of pulse ([γ-$^{32}$P]-ATP) experiments to assess ATP13A3 EP formation (quantified in **Fig. 3L**) and the ATP+SPM-dependent dephosphorylation (quantified in **H**) in the absence or presence of purified AZ2 WT or the ATP13A3-binding-deficient mutant AZ2 Y179A at ATP13A3:AZ2 molar ratios of 1:1 and 1:3. Data are presented as mean ± SEM of *n* = 4 independent biological replicates, with individual data points shown.

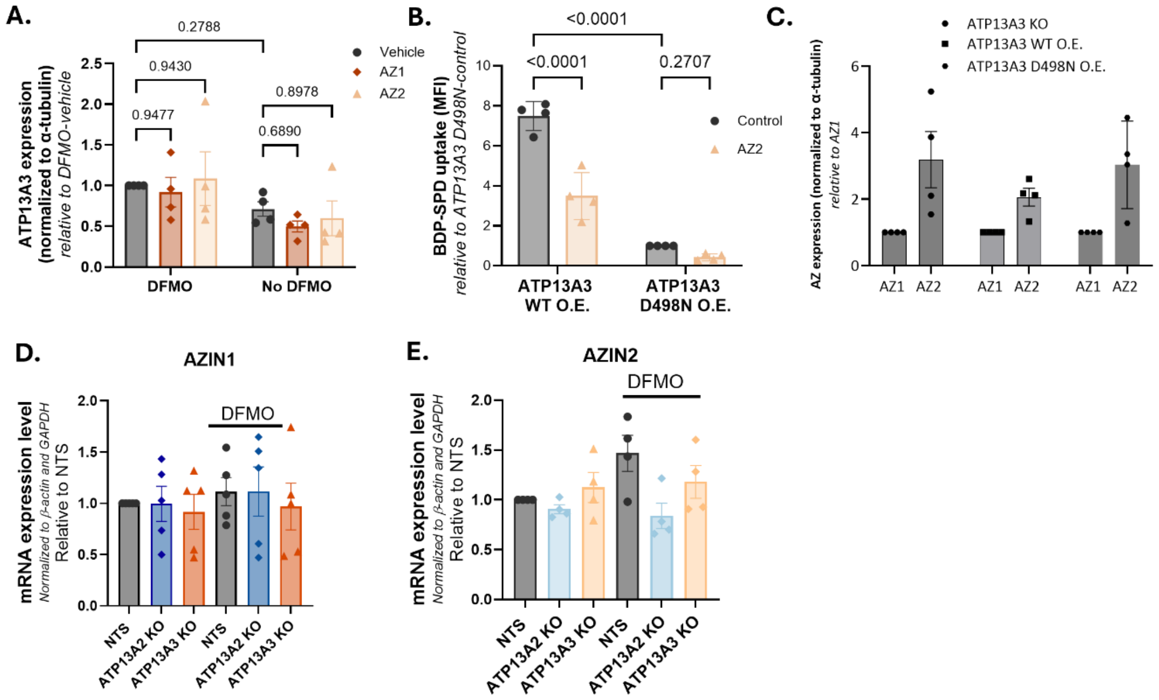


**Figure S9: Effect of AZ1 and AZ2 on ATP13A3 protein expression, AZIN1 and AZIN2 mRNA expression and uptake. (A)** Quantification of ATP13A3 expression in HEK293T NTS cells transfected with Fugene HD (vehicle), AZ1-HA or AZ2-HA in the absence or presence of 20 h pre-treatment with 1.5 mM DFMO. ATP13A3 expression was normalized to α-tubulin. Data are represented as the mean ± SEM of *n* = 4 independent experiments. Two-way ANOVA with Tukey's multiple-comparisons test. **(B)** BODIPY-spermidine (BDP-SPD; 1 µM, 30 min) uptake in parental HEK293T cells stably expressing ATP13A3 WT or ATP13A3 D498N transfected with AZ2-HA. Uptake was normalized to the non-transfected ATP13A3 D498N cells (control). Data are presented as mean ± SEM of *n* = 4 independent biological replicates, with individual data points shown. Two-way ANOVA with Tukey's multiple-comparisons test. **(C)** Quantification of AZ1-HA and AZ2-HA expression in ATP13A3 KO, ATP13A3 WT O.E. and ATP13A3 D498N O.E. cells. AZ expression was normalized to α-tubulin. Data are represented as the mean ± SEM of *n* = 4 independent experiments. **(D-E)** mRNA expression levels of *AZIN1* **(D)**, and *AZIN2* **(E)** in HEK293T NTS and ATP13A3 KO cells in the absence or presence of 24 h pre-treatment with 1.5 mM DFMO. Data are presented as relative expression (2^−ΔΔCt). Ct values were normalized to β-actin (ACTB) and GAPDH, and expression was normalized to NTS. Bars represent mean ± SEM of *n* = 4-5 independent biological replicates, with individual data points shown. Statistical analyses were performed on ΔCt values using one-way ANOVA followed by Dunnett's multiple comparisons test.

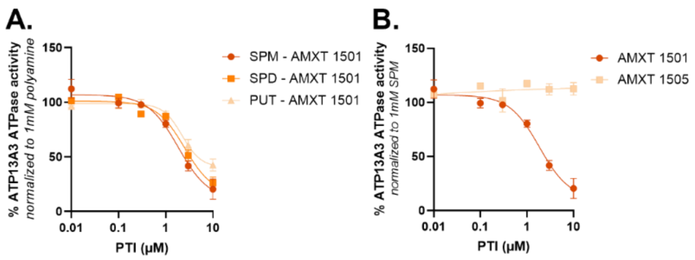


**Figure S10: (A)** Putrescine (PUT), spermidine (SPD), and spermine (SPM)-stimulated ATPase activity of purified ATP13A3 measured in the presence of increasing concentrations of AMXT 1501. ATPase activity was normalized to the activity measured in the presence of 1 mM PUT, SPD or SPM in the absence of AMXT 1501. Data are presented as mean ± SEM of n = 3 independent biological replicates. **(B)** SPM-stimulated ATP13A3 ATPase activity in the presence of increasing concentrations of AMXT 1501 or AMXT 1505. ATPase activity was normalized to the activity measured at 1 mM SPM in the absence of AMXT 1501. Data are presented as mean ± SEM of n = 3 independent biological replicates.

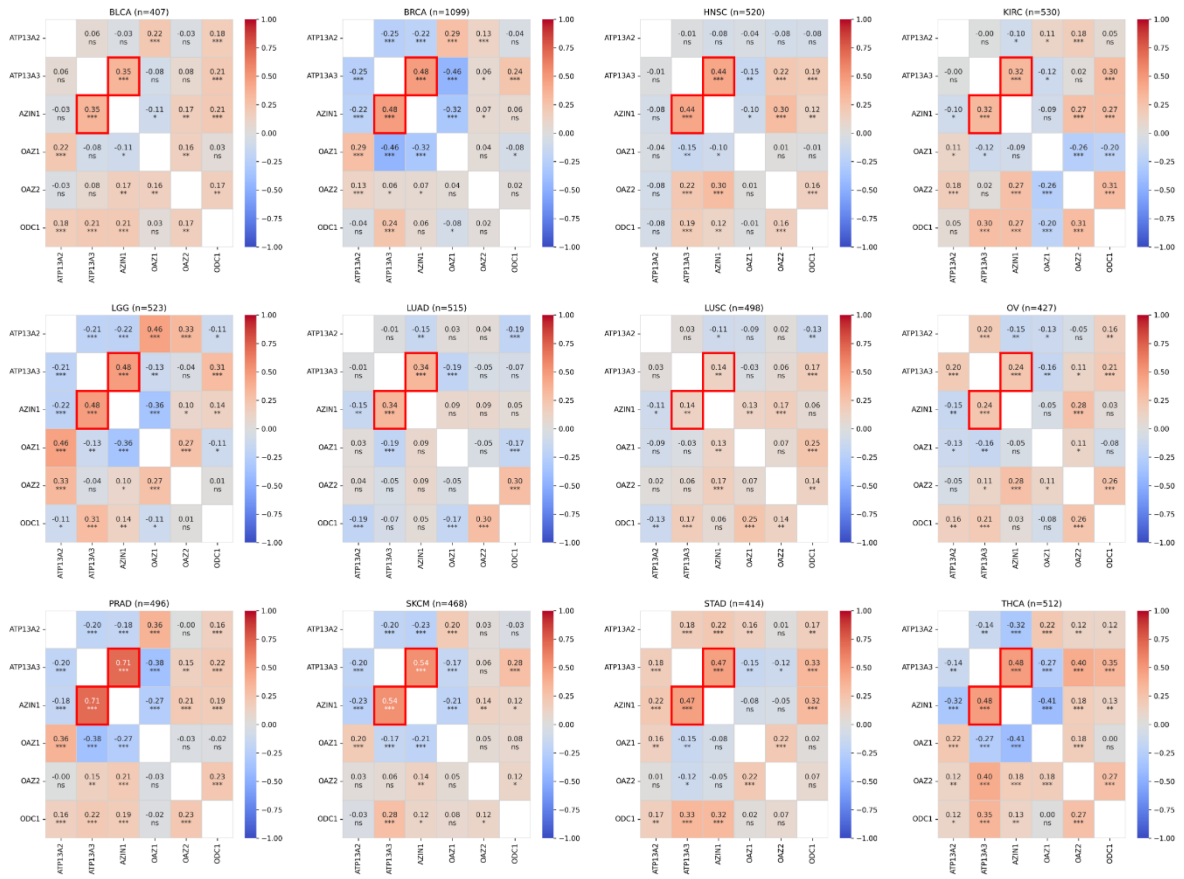


**Figure S11:** Spearman rank correlation (Spearman's ρ) was used to quantify co-expression relationships between all pairs of the 6 polyamine pathway genes (*ATP13A2*, *ATP13A3*, *AZIN1*, *OAZ1*, *OAZ2*, *ODC1*) within tumor samples of each of the 12 TCGA cancer types. The correlation was computed within each cancer type separately, yielding 15 unique gene pairs per cancer type. The correlation between AZIN1 and ATP13A3 was highlighted by the red boxes. Benjamini-Hochberg false discovery rate (BH-FDR) correction was applied globally across all 180 tests. Significance thresholds are indicated in each box: BH-FDR < 0.05 (*); BH-FDR < 0.01 (**); BH-FDR < 0.001 (***) or not significant (ns).

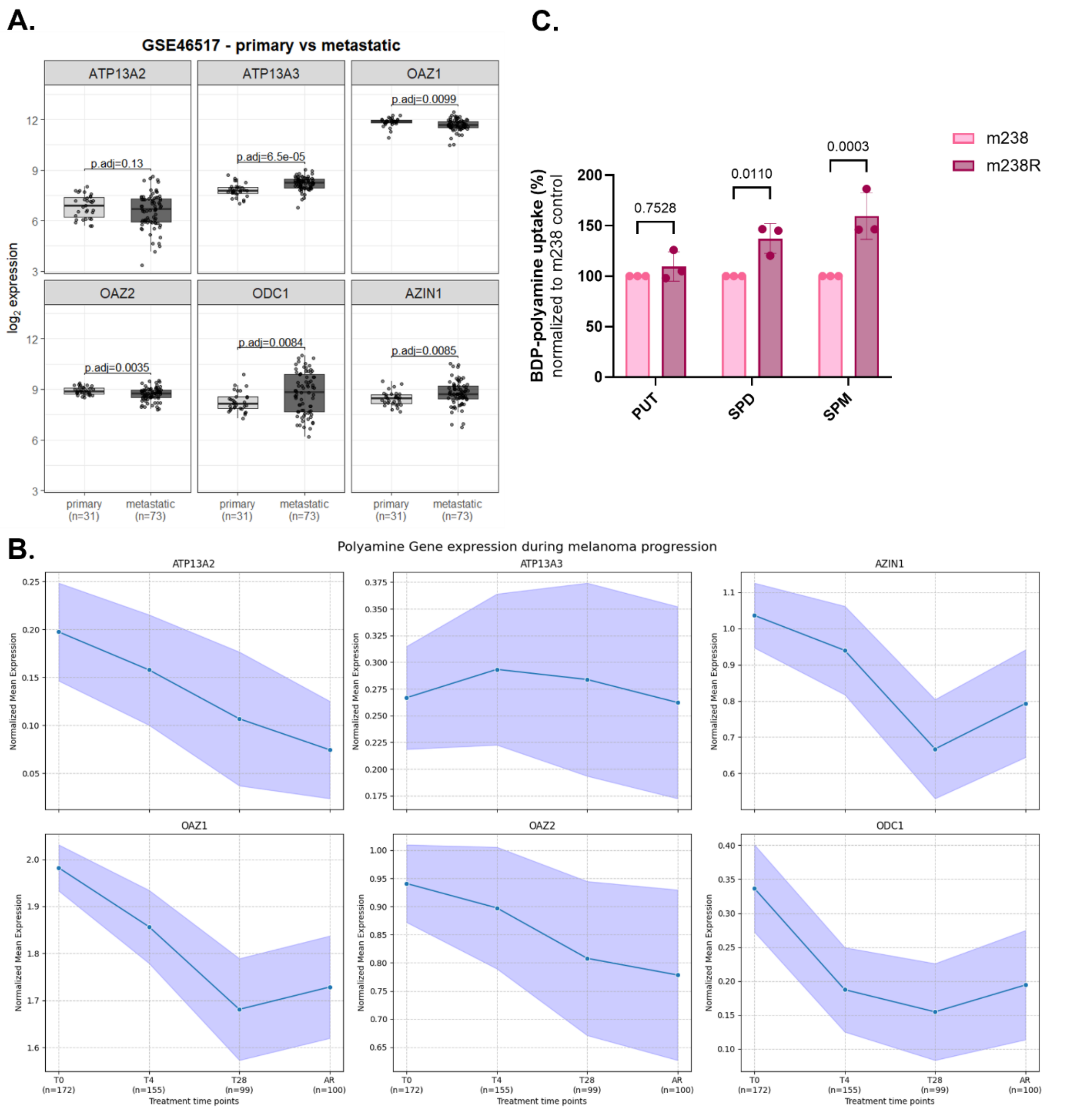


**Figure S12: (A)** The Gene Expression Omnibus (GEO) dataset GSE46517 was used to compare mRNA expression [GEO, $\log_2$ intensity] of *ATP13A2*, *ATP13A3*, *OAZ1–2*, *ODC1* and *AZIN1* between primary (n = 31) and metastatic (n = 73) melanoma tumours. Groups were compared by Welch t-test with Benjamini–Hochberg correction (adjusted p-values shown). **(B)** The Gene Expression Omnibus (GEO) dataset GSE116237 (single-cell RNA-seq of drug-treated melanoma cells: BRAF V600E PDX melanoma (MEL006) cells) was used to calculate the per cell normalized mRNA expression values of the 6 polyamine pathway genes (*ATP13A2*, *ATP13A3*, *AZIN1*, *OAZ1*, *OAZ2*, *ODC1*) at different treatment time points: T0 (pretreatment); T4 = 4 days; T28 = 28 days; AR = acquired resistance at end of treatment. Treatment was a combination of dabrafenib and trametinib. The plots show the normalized mean expression and 95% confidence interval. Differential expression between time points was assessed using the two-sided Mann-Whitney U test (Wilcoxon rank-sum test) with Benjamini-Hochberg correction. Expression levels of *ATP13A2*, *AZIN1*, *OAZ1* and *ODC1* were significantly different adjusted p-values < 0.001 for the comparison T0 – AR. For *ATP13A3* and *OAZ2*, the difference between T0 and AR was marginally significant (adjusted p-values = 0.042 and 0.043) with marginal log2FC for *ATP13A3* (-0.004). **(C)** Uptake of 1 µM BODIPY-polyamines (BDP-PUT, BODIPY-putrescine; BDP-SPD, BODIPY-spermidine; and BDP-SPM, BODIPY-spermine) in melanoma m238 or m238 resistant (m238R) cells. Cells were treated for 30 min with BDP-polyamines in the presence of 1 mM aminoguanidine. Uptake was quantified as mean fluorescence intensity and normalized to m238 cells. Data are presented as mean ± SEM of n= 3 independent biological replicates, with individual data points shown. Two-way ANOVA with Sidak's multiple comparisons test.
